# Protein subcellular relocalization and function of duplicated flagellar calcium binding protein genes in honey bee trypanosomatid parasite

**DOI:** 10.1101/2023.06.17.545447

**Authors:** Xuye Yuan, Tatsuhiko Kadowaki

## Abstract

The honey bee trypanosomatid parasite, *Lotmaria passim*, contains two genes that encode the flagellar calcium binding protein (FCaBP) through tandem duplication in its genome. FCaBPs localize in the flagellum and cell body of *L. passim* through specific N-terminal sorting sequences. This finding suggests that this is an example of protein subcellular relocalization resulting from gene duplication, altering the intracellular localization of FCaBP. However, this phenomenon may not have occurred in *Leishmania*, as one or both of the duplicated genes have become pseudogenes. Multiple copies of the *FCaBP* gene are present in several *Trypanosoma* species and *Leptomonas pyrrhocoris*, indicating rapid evolution of this gene in trypanosomatid parasites. The N-terminal flagellar sorting sequence of *L. passim* FCaBP1 interacts with the BBSome complex, while those of *Trypanosoma brucei* and *Leishmania donovani* FCaBPs do not direct GFP to the flagellum in *L. passim*. These results suggest that the N-terminal flagellar sorting sequence of FCaBP1 has co-evolved with the BBSome complex in each trypanosomatid species. Deletion of the two *FCaBP* genes in *L. passim* affected growth and impaired flagellar morphogenesis and motility, but it did not impact host infection. Therefore, *FCaBP* represents a duplicated gene with a rapid evolutionary history that is essential for flagellar structure and function in a trypanosomatid parasite.

## Introduction

Gene duplication is considered the main source of new genes (Innan and Kondrashov, 2010; Ohno, 1970; Zhang, 2003; Kuzmin, Taylor, and Boone, 2022). However, to maintain duplicate genes in a genome, functional divergence is usually necessary, with a few exceptions. Functional divergence can occur through neofunctionalization, where one of the duplicates develops a new function while the other retains the original function of the ancestral gene (Ohno, 1970). Alternatively, subfunctionalization can take place, where the ancestral functions are divided between the duplicate genes. For example, their combined levels or patterns of activity could be equivalent to the original single gene (Hughes, 1994; Force et al., 1999; Stoltzfus, 1999). These types of functional divergence can modify the function of the encoded protein or the gene’s expression pattern. Subcellular localization is also crucial for a protein’s function within a cell. Protein subcellular relocalization (PSR) has been proposed as a mechanism for the functional divergence and retention of duplicate genes (Byun-McKay and Geeta, 2007; Marques et al., 2008). PSR allows for the rapid subdivision of ancestral localization or acquisition of new localization following gene duplication. After neolocalization or sublocalization, the expression pattern and function of the genes can further diverge. Although this idea was questioned by comparing the frequency of PSR between singletons and duplicates (Qian and Zhang, 2009), there are numerous examples of PSR for duplicated genes (Wang et al., 2009; Byun and Singh, 2013; Liu, Pan, and Adams, 2014).

The flagellar calcium binding protein (FCaBP or Calflagin) was discovered as one of the abundant proteins in the flagellar membrane of *Trypanosoma brucei* and *Trypanosoma cruzi*. It contains four EF-hand calcium binding domains and is considered a conserved signaling protein among trypanosomatid parasites (Buchanan et al., 2005; Wingard et al., 2008). FCaBP is acylated with myristate and palmitate at the N-terminus, which is crucial for its localization in the flagellum and association with lipid raft microdomains (Godsel and Engman, 1999; Emmer et al., 2009; Tyler et al., 2009). Additionally, the N-terminal amino acid sequence of FCaBP is necessary for its targeting to the flagellum in *T. cruzi* (Maric et al., 2011). Although the phenotypes of knocking down FCaBPs in *T. brucei* have been reported (Emmer et al., 2010), the physiological functions of FCaBPs are not well understood. Proteins in the flagellum (and cilium) are initially synthesized in the cell body and then transported to the flagellum through intraflagellar transport (IFT). IFT requires IFT complexes (IFT-A and IFT-B) and motor proteins, including kinesin-2 for anterograde transport and IFT-dynein for retrograde transport, along the axonemal microtubules (Cole et al., 1998; Pazour, Dickert, and Witman, 1999; Piperno and Mead, 1997). The IFT complexes act as carriers, and BBSome and tubby-like protein (TULP) function as adapters by binding to specific sets of flagellar membrane proteins (Blacque et al., 2004; Lechtreck et al., 2009; Nachury et al., 2007; Singh et al., 2020; Badgandi et al., 2017; Legué and Liem, 2019). For example, BBSome is necessary for the exit of a dually acylated phospholipase D from cilia in *Chlamydomonas* (Lechtreck et al., 2009). Therefore, FCaBP likely requires IFT complexes and either BBSome or TULP for its import and export to the flagellum.

*Lotmaria passim* is the most prevalent trypanosomatid parasite that infects honey bees worldwide (Regan et al., 2018; Stevanovic et al., 2016; Arismendi et al., 2016; Ravoet et al., 2015). It specifically colonizes the hindgut of honey bees, affecting the host’s physiology, and it may be associated with winter colony loss (Ravoet et al., 2013; Liu et al., 2020). *L. passim* is a monoxenous parasite that only infects bees and can serve as a model to study the roles of flagellar formation and function for infecting the various hosts of trypanosomatid parasites.

In this study, we identified two *FCaBP* genes in the genomes of *L. passim* and closely related species, resulting from tandem duplication. By analyzing the gene in other trypanosomatid species’ genomes, the protein subcellular localizations, and N-terminal sorting sequences, we have uncovered their evolutionary history in relation to the flagellar protein transport machinery. Deletion of these genes in *L. passim* has also revealed novel physiological functions of FCaBPs. Therefore, *FCaBP* serves as an example to understand how duplicated genes have diversified during the evolution of trypanosomatid parasites.

## Results

### *L. passim* contains two FCaBPs with flagellar and cell body localizations

The genome of *L. passim* encodes two *FCaBP* genes (*LpFCaBP1* and *LpFCaBP2*) that are separated by 1063 bp, suggesting tandem gene duplication. To determine their localizations, we expressed GFP-tagged versions of LpFCaBP1 and LpFCaBP2 (at C-terminus) in *L. passim*. LpFCaBP1-GFP was highly enriched in the flagellum, similar to FCaBPs in *T. cruzi* and *T. brucei* (Godsel and Engman, 1999; Emmer *et al*., 2009). On the other hand, most of LpFCaBP2-GFP was present at the plasma membrane of the cell body (Fig. 1). We also observed that GFP alone is uniformly distributed in the cell body, while the paraflagellar rod protein 5 (LpPFRP5)-GFP fusion protein is specific to the flagellum (Fig. 1). These findings indicate that the two duplicated *FCaBP* genes encode proteins with distinct subcellular localizations.

**Figure 1.**
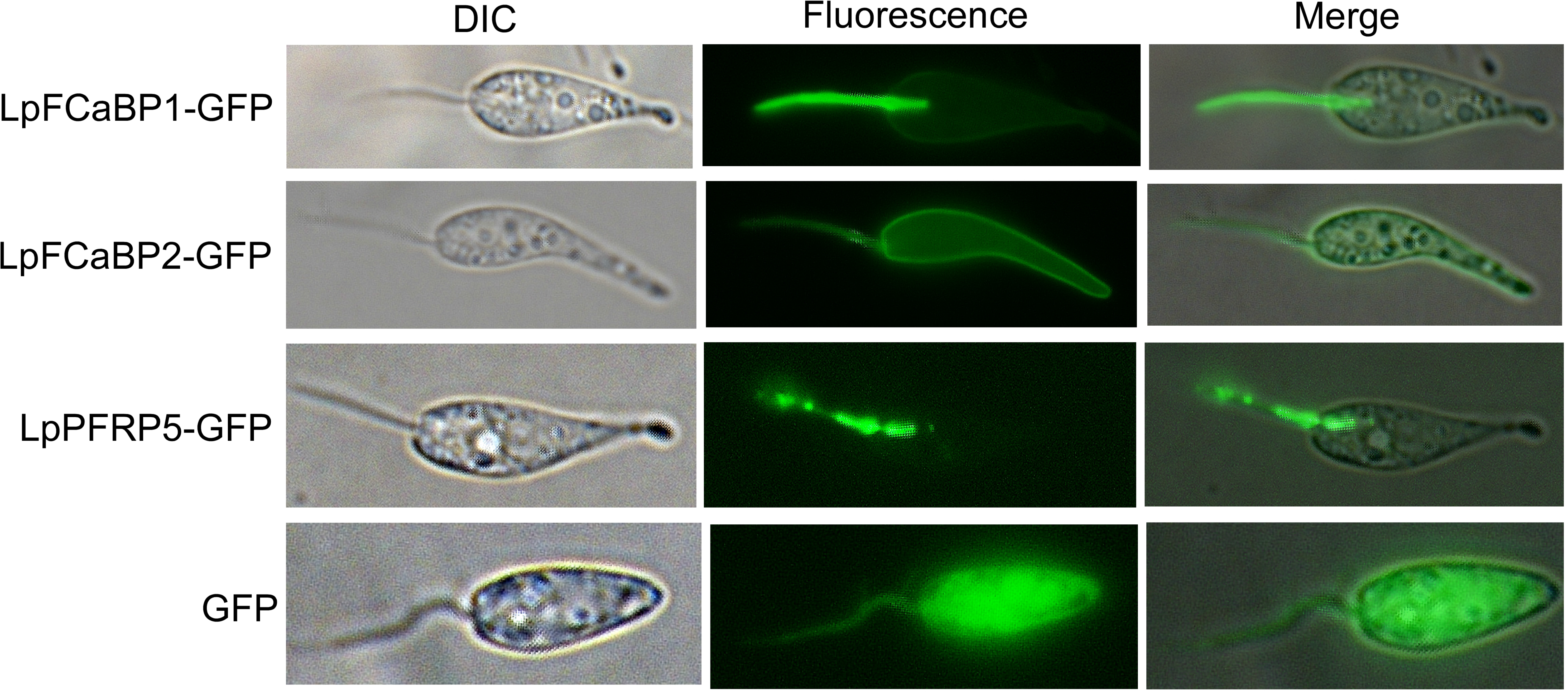
Subcellular localization of LpFCaBP1- and 2-GFP fusion proteins. *Lotmaria passim* expressing LpFCaBP1-GFP, LpFCaBP2-GFP, LpPFRP5-GFP, or GFP were examined under visible (DIC) or fluorescence (Fluorescence) light. Merged images are also shown. The flagellum is located at the anterior side of the parasite.

### The N-terminal 16 amino acids of LpFCaBP function as either flagellum or cell body sorting sequence

By aligning the amino acid sequences of LpFCaBP1 and LpFCaBP2, we found that they are almost identical except for their N-terminal ends (Supplementary file 1). To determine the flagellar and cell body sorting sequences, we swapped the N-terminal 16 or 28 amino acids between LpFCaBP1 and LpFCaBP2 in GFP fusion proteins. The localization of the chimeric proteins was directed by the N-terminal 16 and 28 amino acids (Fig. 2), while the remaining amino acids, which constitute the EF-hand calcium binding domains, did not play a critical role. The N-terminal 16 amino acids of LpFCaBP1 and LpFCaBP2 were sufficient to direct GFP to the flagellum and cell body, respectively (Fig. 2), indicating that they represent the flagellar and cell body sorting sequences.

**Figure 2.**
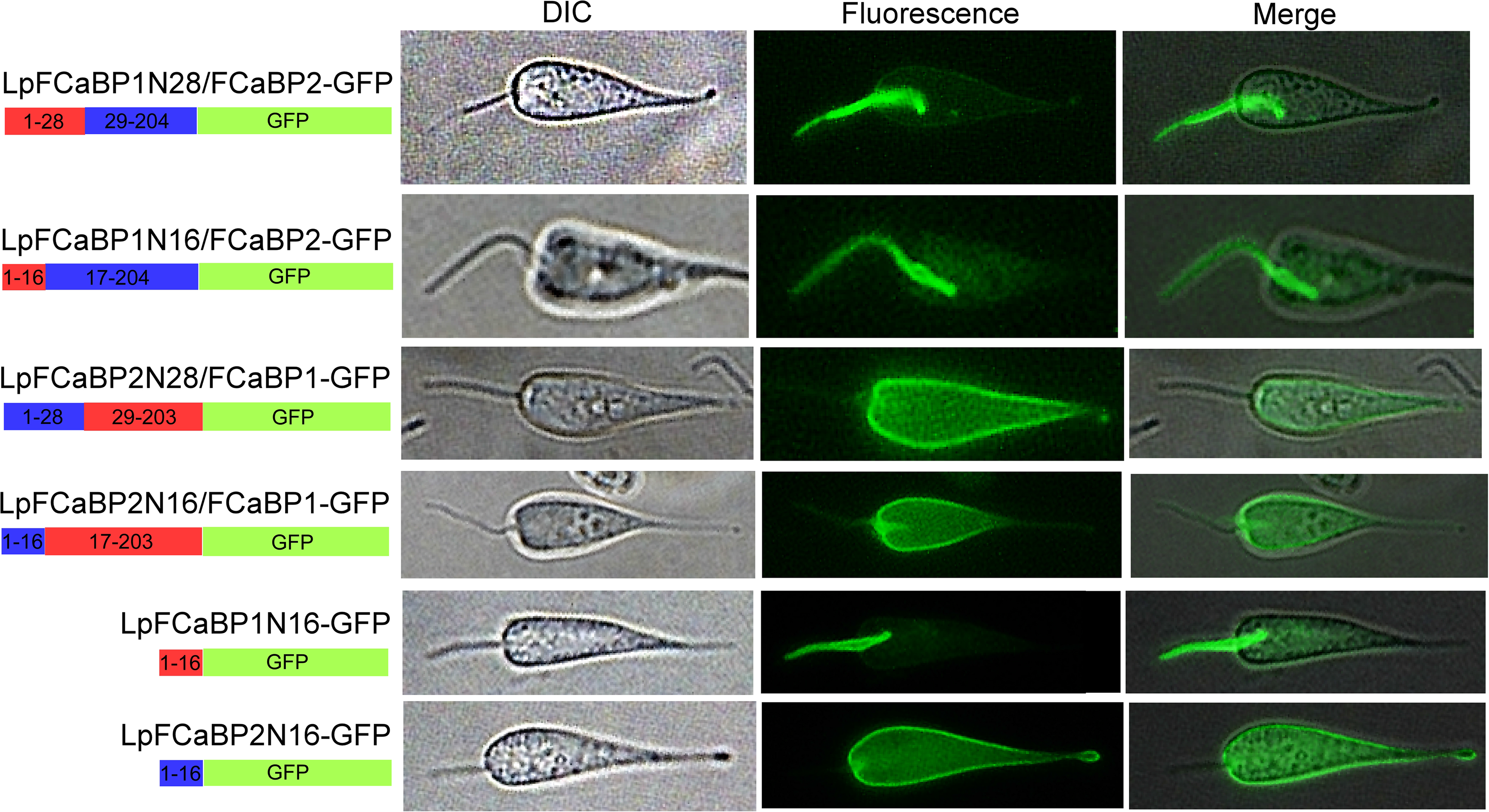
Flagellum and cell body sorting sequences of LpFCaBP1 and 2. The N-terminal 16 or 28 amino acids of LpFCaBP1 (in red) and LpFCaBP2 (in blue) were swapped as illustrated in the diagram. The positions of amino acids in the cores of LpFCaBP1 and 2 proteins are also indicated. GFPs fused with the N-terminal 16 amino acids of LpFCaBP1 and 2 are shown at the bottom.

### *FCaBP* genes have rapidly evolved in trypanosomatid parasites

We examined *FCaBP* genes in various trypanosomatid parasites’ genomes. *T. brucei* has four *FCaBP* and the related genes as previously reported (Emmer *et al*., 2010). *T. cruzi* appears to contain the large number of *FCaBP* genes due to its repetitive genome (Arner *et al*., 2007). *Trypanosoma theileri* genome has three *FCaBP* genes (Supplementary file 2). These results suggest that *FCaBP* gene duplicated in the common ancestor of *Trypanosoma* and Leishmaniinae and has undergone further duplication in each *Trypanosoma* species.

In the genomes of various Leishmaniinae species, microsynteny around *FCaBP* genes is well conserved with the same orders and orientations of genes encoding three novel proteins, dynein light chain, and sucrose-phosphate synthase-like protein (Fig. 3). In *Leishmania infantum*, *Leishmania braziliensis*, *Leishmania mexicana*, and *Leishmania donovani,* two *FCaBP* genes are present but *FCaBP2* appears to have become a pseudogene by generating a stop codon in 5’ end of the ORF (open reading frame). If this gene is translated, the truncated protein without N-terminal sorting sequence is synthesized. Thus, above four *Leishmania* species contain only *FCaBP1*. Intriguingly, two *FCaBP* genes in *Leishmania major* have become pseudogenes by accumulating multiple stop codons in the ORFs (Fig. 3).

**Figure 3.**
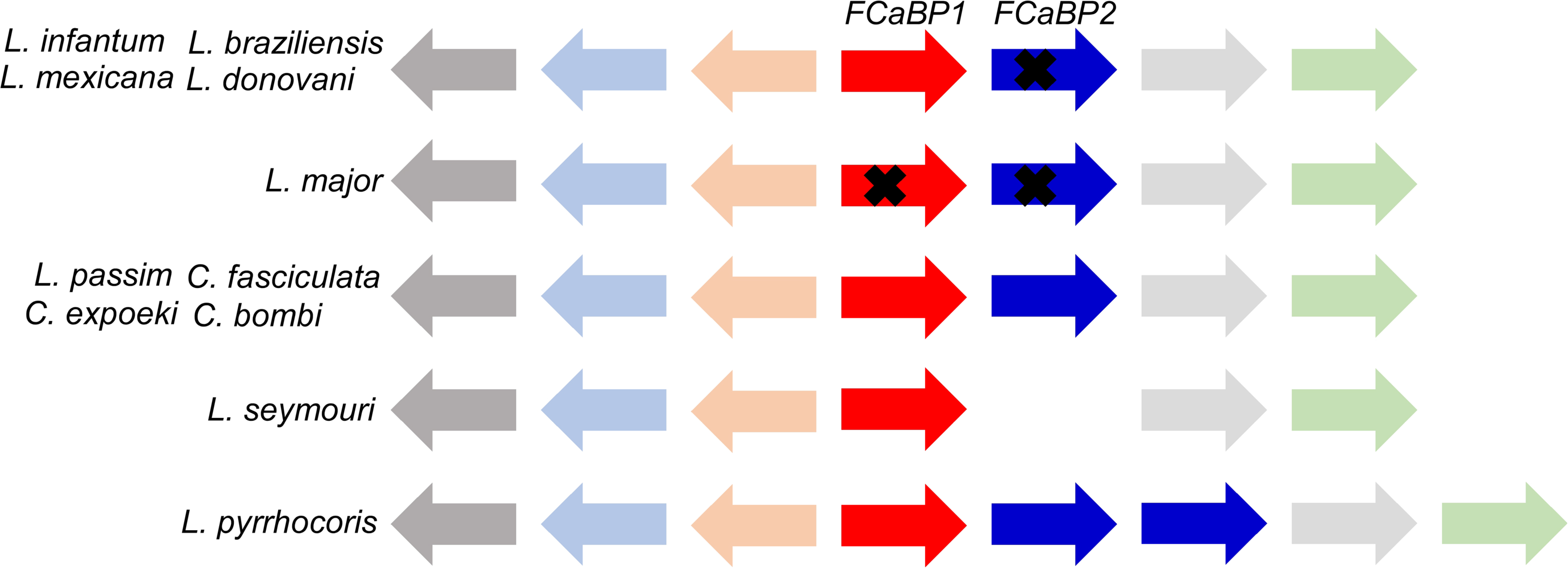
Conserved microsynteny of *FCaBP* genes in Leishmaniinae species. The microsynteny containing *FCaBP* genes is highly conserved in *Leishmania infantum, Leishmania braziliensis, Leishmania mexicana, Leishmania donovani, Leishmania major, Lotmaria passim, Crithidia fasciculata, Crithidia expoeki, Crithidia bombi, Leptomonas seymouri*, and *Leptomonas pyrrhocoris*. *FCaBP2* is a pseudogene (indicated by ×) in *L. infantum, L. braziliensis, L. mexicana*, and *L. donovani,* while both *FCaBP1* and *2* are pseudogenes in *L. major*. *FCaBP* genes are flanked by three novel genes (beige, light grey, and light green arrows), as well as dynein light chain (light blue arrow) and sucrose-phosphate synthase-like protein (dark grey arrow). The orientation of each gene is shown.

In species related to *L. passim* (Schwarz et al., 2015), *Crithidia fasciculata*, *Crithidia expoeki*, and *Crithidia bombi* have two *FCaBP* genes, while *Leptomonas seymouri* has only *FCaBP1* gene (Fig. 3). *Leptomonas pyrrhocoris* contains *FCaBP1* and two *FCaBP2* genes based on the comparison of N-terminal sorting sequences (Supplementary file 1). These results demonstrate that *FCaBP* gene has rapidly evolved in trypanosomatid parasites.

### Flagellar localization of FCaBP depends on both the sorting sequence and trans-acting factors in *L. passim*

To test the conservation and plasticity of the flagellar and cell body sorting sequences of FCaBPs, we expressed GFP fusion proteins containing N-terminal sorting sequences from various trypanosomatid parasites in *L. passim*. Figure 4A shows the N-terminal ends of aligned FCaBPs from *T. cruzi, T. brucei, L. donovani*, C. *fasciculata, L. seymouri,* and *L. passim.* Glycine and cysteine residues targeted for myristoylation and palmitoylation are well conserved; however, there are more variations in the sorting sequences of FCaBPs compared to the main sequences with EF-hand calcium binding domains. Because TcFCaBP and TbFCaBP (Calflagin, Tb24) are flagellar proteins, their N-terminal amino acids should function as flagellar sorting sequences in *T. cruzi* and *T. brucei*, respectively (Emmer *et al*., 2009; Maric *et al*., 2011). However, we found that GFP fused with the sorting sequence of either TbFCaBP or LdFCaBP1 is localized in the cell body of *L. passim*. GFP with the sorting sequence of TcFCaBP is present in both cell body and the proximal part of flagellum (Fig. 4B). The sorting sequences of CfFCaBP1 and 2 directed GFP to both cell body as well as flagellum and cell body, respectively. GFP with the sorting sequence of LsFCaBP1 is primarily localized in flagellum (Fig. 4B). These results suggest that dually acylated FCaBP is targeted to the cell body membrane by default, and efficient flagellar localization requires trans-acting factor(s) to recognize the specific flagellar sorting sequence.

**Figure 4.**
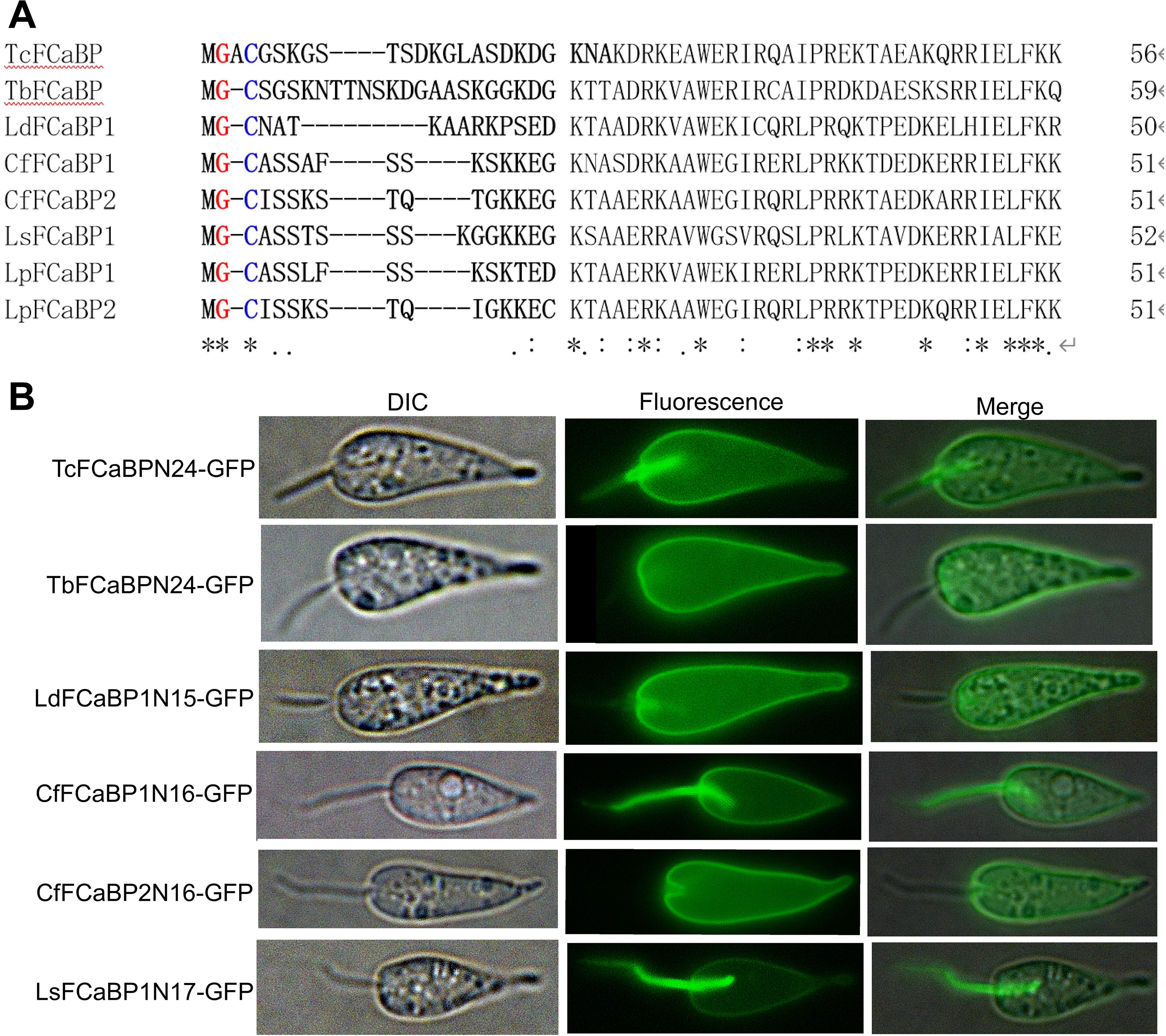
Subcellular localization of GFPs with N-terminal sorting sequences of FCaBPs from various trypanosomatid parasites. (A) N-terminal parts of the aligned sequences of *Trypanosoma cruzi, Trypanosoma brucei, Leishmania donovani, Crithidia fasciculata, Leptomonas seymouri*, and *Lotmaria passim* FCaBPs are shown. The amino acids fused to GFP in (B) are indicated in bold letters. Glycine (red) and cysteine (blue) residues are targeted for myristoylation and palmitoylation, respectively. Conserved amino acids are shown by asterisks, and similar amino acids are indicated by either a full stop (.) or a colon (:). (B) Subcellular localization of GFPs fused with the N-terminal sorting sequences in (A).

### The N-terminal sorting sequence of LpFCaBP1 interacts with the BBSome complex but not TULP

Since the import and export of flagellar membrane proteins depend on either the BBSome complex or TULP (Lechtreck, 2022), we first tested the interaction between GFP fused with N-terminal flagellar sorting sequence of LpFCaBP1 (LpFCaBP1N16-GFP) and Flag-tagged LpBBS1 or LpTULP in the parasites expressing both proteins by immunoprecipitation. However, we were unable to detect binding with either protein. We thus used an engineered biotin ligase, ultraID (Kubitz *et al*., 2022), instead of GFP. This fusion protein allowed us to identify the interacting proteins through biotinylation. In *L. passim* expressing both LpFCaBP1N16-ultraID and Flag-tagged LpBBS1 or LpTULP, ultraID was detected in the proximal part of the flagellum (Fig. 5A). Among the biotinylated proteins captured by streptavidin-agarose, Flag-LpBBS1, but not LpTULP, was detected (Fig. 5B). The apparent size of Flag-LpTULP was larger than that of Flag-LpBBS1 and the expected size (59 kDa) (Fig. 5B). LpTULP may have post-translational modification or migrate slowly through 10 % SDS-PAGE. These findings suggest that the flagellar transport of LpFCaBP1 is likely mediated by the BBSome complex.

**Figure 5.**
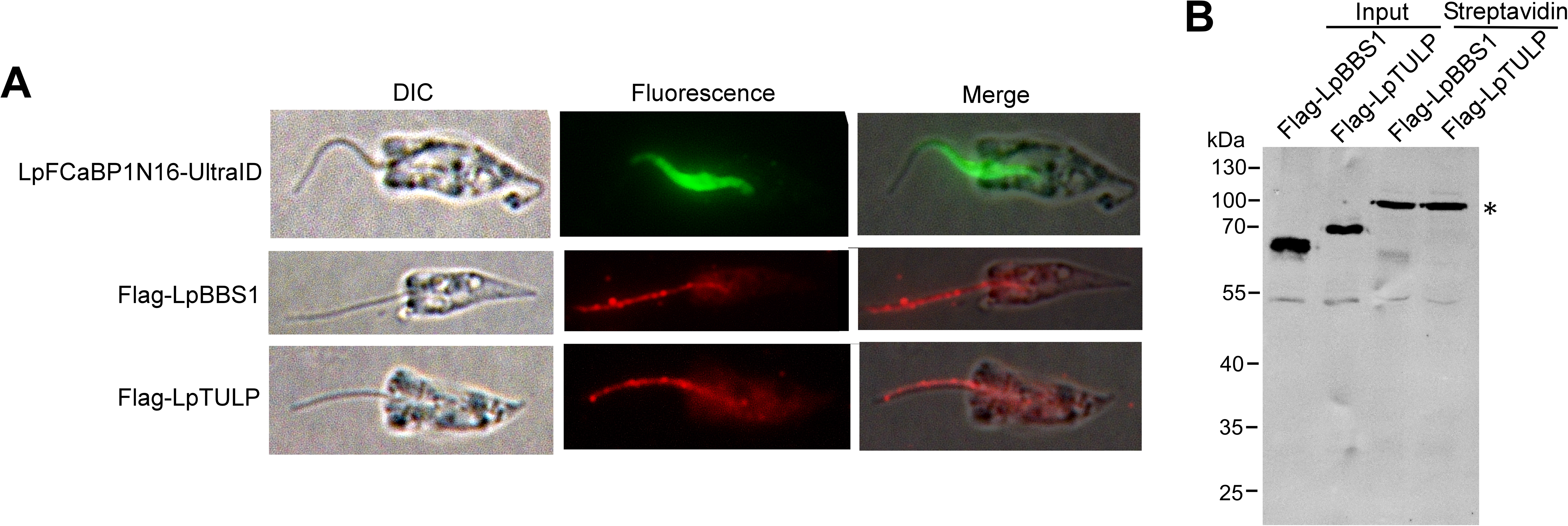
Interaction of LpFCaBP1 N-terminal sorting sequence with LpBBS1. (A) Subcellular localization of ultraID with the N-terminal sorting sequence of LpFCaBP1 (LpFCaBP1N16-ultraID), Flag-tagged LpBBS1 (Flag-LpBBS1), and LpTULP (Flag-LpTULP) by immunofluorescence. (B) Lysates of parasites expressing both LpFCaBP1N16-ultraID and Flag-LpBBS1 or Flag-LpTULP were subjected to immunoprecipitation using Flag Fab-Trap Agarose (Input) or streptavidin agarose precipitation (Streptavidin) to capture biotinylated proteins. The precipitates were analyzed by western blot using an anti-Flag tag antibody. An asterisk indicates a major biotinylated 100 kDa band that non-specifically cross-reacted with the antibody.

### LpFCaBPs are essential for flagellar morphogenesis and motility but not the host infection of *L. passim*

To determine the functions of LpFCaBPs, we deleted both *LpFCaBP1* and *LpFCaBP2* genes in *L. passim* by replacing them with hygromycin resistance gene by CRISPR (Fig. 6A). Although LpFCaBP1 and 2 primarily localize in the flagellum and cell body, they could be still functionally redundant. By genomic PCR, we identified both heterozygous (+/-) and homozygous (-/-) deletion clones of *FCaBPs* (Fig. 6B). To confirm the disruption of genes, we tested *LpFCaBP1* and *2* mRNA expression in wild type, heterozygous and homozygous deletion parasites by RT-PCR using the gene-specific reverse primer. As shown in Figure 6C, both mRNAs are absent in the homozygous deleted parasites (*LpFCaBP1/2-/-*).

**Figure 6.**
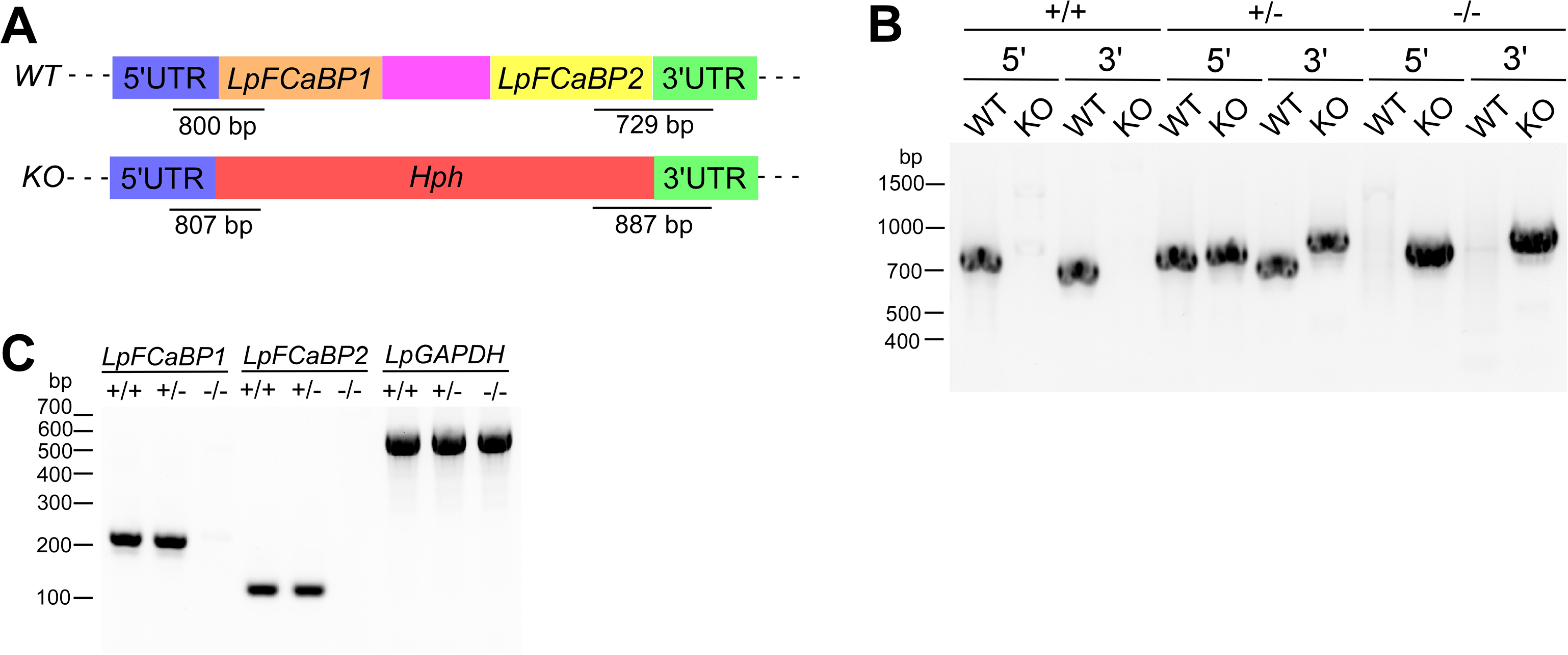
Deletion of *LpFCaBP1* and *2* genes by CRISPR. (A) Schematic representation of wild-type (*WT*) and deleted (*KO*) alleles of *LpFCaBPs* generated by CRISPR/Cas9-induced homology-directed repair. 5’ and 3’ untranslated regions (UTRs), open reading frames (ORFs), and the internal spacer of *FCaBP1* and *2*, hygromycin resistance gene *(Hph)*, are shown in blue, green, orange, yellow, purple, and red, respectively. The expected sizes of PCR products to detect *WT* and *KO* alleles (not to scale) are also shown. (B) Genomic DNAs of wild-type *L. passim* (+/+), heterozygous (+/-), and homozygous (-/-) mutants of *LpFCaBPs* were analyzed by PCR to detect *5’WT, 5’KO, 3’WT,* and *3’KO* alleles. Sizes of the DNA molecular weight markers are shown on the left. (C) Detection of *LpFCaBP1* and *2*, as well as *LpGAPDH* mRNAs, in *LpFCaBPs* heterozygous (+/-) and homozygous (-/-) mutants together with wild-type *L. passim* (+/+) by RT-PCR. Sizes of the DNA molecular weight markers are shown on the left.

Compared to wild-type parasites, *LpFCaBP1/2-/-* grew faster during the logarithmic phase but reached the stationary phase at a lower density (Fig. 7A and B). *LpFCaBP1/2-/-* parasites are viable but have the distinct growth characteristics. Wild-type parasites extend the flagella and start actively moving at the early stage of stationary phase of culture (at 60 h after the culture). We found that *LpFCaBP1/2-/-* parasites have shorter flagella and less active movement compared to wild-type (Fig. 7C, D, E, and Supplementary videos 1 and 2). These results indicate that *LpFCaBP1* and *2* genes are necessary for flagellar morphogenesis as well as motility of *L. passim*. At the late stage of stationary phase of culture (at 96 h after the culture), wild-type parasites form rosettes (clusters of cells with their flagella toward the center of the cluster). However, they were absent with *LpFCaBP1/2-/-* parasites which remain as individuals without clustering at the bottom of culture plate (Fig. 7F). We then compared the infection of wild-type and *LpFCaBP1/2-/-* parasites in honey bee gut and found that they infect the host at the comparable level (Fig. 7G). Thus, less motile *L. passim* with short flagellum by the loss-of function of *LpFCaBP* genes is still capable of infecting honey bee hindgut effectively.

**Figure 7.**
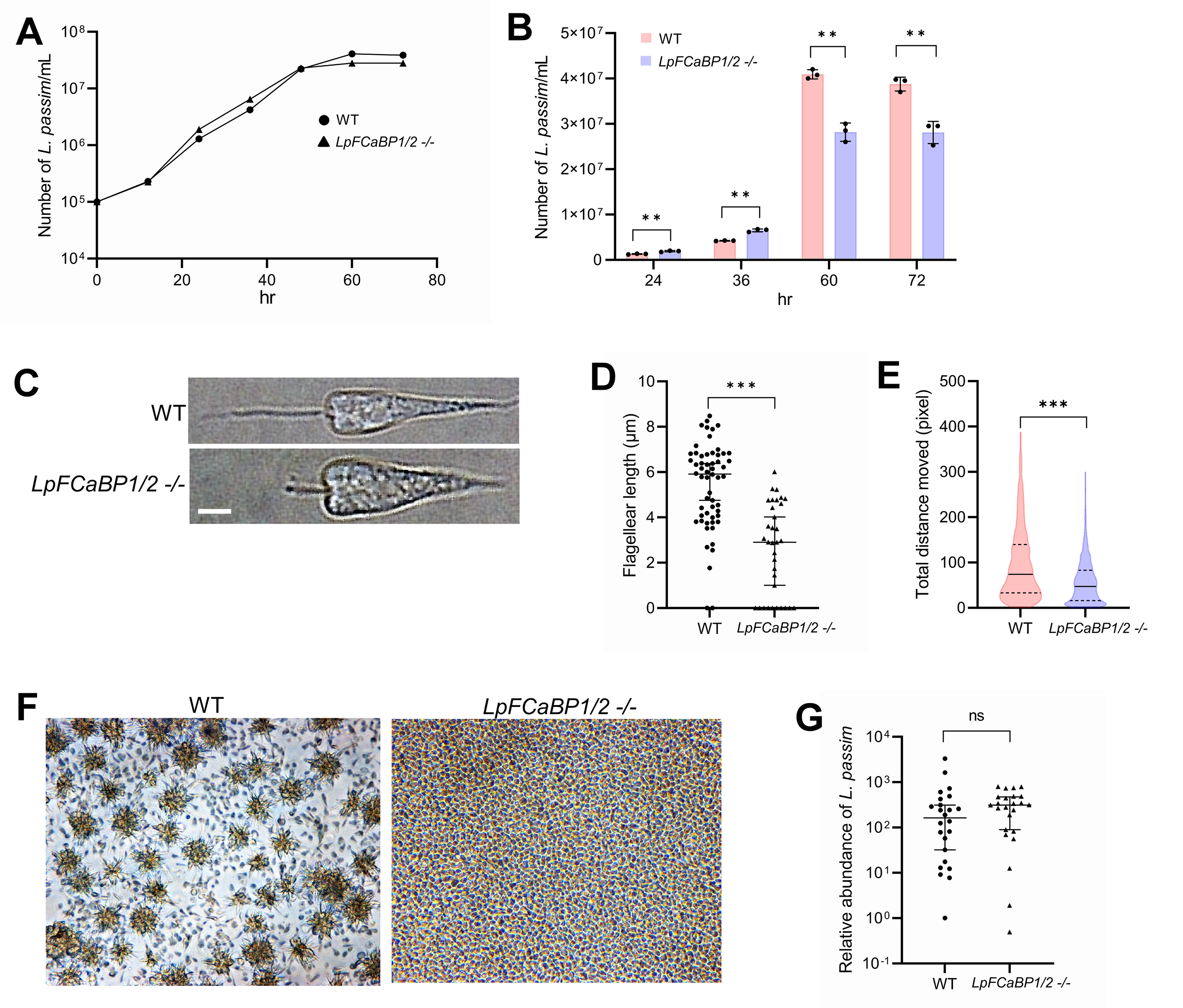
Phenotypes of *LpFCaBPs* deleted mutant under culture conditions and in honey bees. (A) Growth rates of wild-type (WT, circle) and *LpFCaBPs* homozygous mutant (*LpFCaBP1/2 -/-*, triangle) *L. passim* in the modified FP-FB medium were monitored at 28 °C for three days (n = 3). (B) Density of WT and *LpFCaBP1/2 -/-* parasites compared at 24, 36, 60, and 72 hours after culture. Statistical comparisons were carried out using Welch’s *t*-test (_**_). *P* < 0.0076, *P* < 0.005, *P* < 0.0025, and *P* < 0.0056 for 24, 36, 60, and 70 hours, respectively. (C) Morphology of WT and *LpFCaBP1/2 -/-* parasites. Scale bar: 2 μm. (D) Flagellar length of individual WT (n = 58) and *LpFCaBP1/2 -/-* (n = 34) parasites. Statistical comparison was carried out using the Brunner-Munzel test (_***_). *P* < 0.001. (E) Motility (total distance moved for 1 minute) of individual WT (n = 773) and *LpFCaBP1/2 -/-* (n = 2096) parasites shown by a violin plot. Median, as well as the first and third quartiles, are indicated by solid and dashed lines, respectively. Statistical comparison was carried out using the Brunner-Munzel test (_***_). *P* < 0.001. (F) Images of WT and *LpFCaBP1/2 -/-* parasites at 96 hours after culture. The clusters of parasites in WT represent rosettes. (G) The relative abundance of *L. passim* in individual honey bees (n = 24) at 14 days after the infection compared between WT and *LpFCaBP1/2 -/-* parasites. One sample infected by the WT parasite was set as 1, and the median with 95% CI is shown. The Brunner-Munzel test was used for statistical analysis. ns: not significant.

## Discussion

### Rapid evolution of *FCaBP* genes in trypanosomatid parasites

We characterized *FCaBP* genes in the genomes of trypanosomatid parasites and observed that each species, except *L. seymouri*, has more than two *FCaBP* genes in the same genomic contigs (Fig. 3). This suggests that gene duplication occurred in the common ancestor of *Trypanosoma* and Leishmaniinae. While *T. brucei, T. theileri*, and *L. pyrrhocoris* have further duplicated *FCaBP* genes, four *Leishmania* species and *L. seymouri* have lost *FCaBP2* by pseudogenization and deletion, respectively. When we expressed GFPs fused with the sorting sequences of LdFCaBP1 and TbFCaBP in *L. passim*, we observed their localization in the cell body. Meanwhile, GFPs fused with the sorting sequences of TcFCaBP and CfFCaBP1 were present in both the flagellum and cell body (Fig. 4B). These findings indicate that the BBSome complex in *L. passim* may not recognize the flagellar sorting sequences of TbFCaBP and LdFCaBP1, but partially recognizes those of TcFCaBP and CfFCaBP1. Therefore, it appears that dually acylated FCaBP is naturally targeted to the plasma membrane of the cell body, and specific N-terminal sorting sequences and the BBSome complex are required for flagellar sorting. Additionally, the inefficient transport of GFP to the distal part of the flagellum using the TcFCaBP sorting sequence suggests a functional or structural separation in the flagellum (See also Fig. 5A). The lack of functional FCaBP in *L. major* demonstrates that it is not essential for completing the life cycle in sand fly and mammalian hosts, suggesting the presence of other proteins that complement the absence of FCaBP in *L. major*.

In trypanosomatid parasites with two *FCaBP* genes, mutations have accumulated in the 5’ end of the ORF, modifying the N-terminal amino acid sequences after gene duplication. As a result, FCaBP1 and FCaBP2 have different subcellular localizations at the flagellum and cell body, respectively. Thus, this can be considered as one example of PSR to retain duplicated genes. However, this process may not have occurred in the five *Leishmania* species and *L. seymouri*. Without an outgroup paralogous gene or a species with a single *FCaBP* gene, it is difficult to conclude whether this represents neolocalization or sublocalization. However, sublocalization is unlikely due to the absence of *FCaBP2* in the aforementioned species. Since neolocalization to the flagellum requires co-evolution with the BBSome complex, the chance of this occurring is quite low. Therefore, we like to propose that neolocalization to the cell body has occurred, and FCaBP2 in the plasma membrane of the cell body must provide some advantages for *C. fasciculata, L. passim, C. bombi, C. expoeki*, and *L. pyrrhocoris*.

### Interaction between LpFCaBP1 and the BBSome complex in *L. passim*

The flagellar localization of LpFCaBP1 may depend on the interaction between the N-terminal sorting sequence and the BBSome complex, rather than TULP (Fig. 5). These results are consistent with the requirement of the BBSome complex for the exit of dually acylated phospholipase D to the cilia in *Chlamydomonas* (Liu and Lechtreck, 2018). However, the fraction of LpBBS1 interacting with LpFCaBP1N16-ultraID appears to be very small (Fig. 5), possibly due to the limited localization of LpFCaBP1N16-ultraID to the proximal part of the flagellum. It remains to be determined whether the entry/exit of LpFCaBP1 to the flagellum requires the BBSome and IFT complexes.

### Roles of FCaBPs in flagellar morphogenesis and motility of *L. passim*

We found that LpFCaBPs are not essential for the viability of *L. passim*, which is consistent with the lack of FCaBP in *L. major*. However, the loss of LpFCaBPs altered the growth characteristics of *L. passim* in the culture medium and impaired flagellar morphogenesis and motility. The formation of rosettes, which requires the clustering of parasites through flagellar movement (Iovannisci, Plested and Moe, 2010), was also defective in *LpFCaBPs-*deleted parasites (Fig. 7). In contrast, knocking down *TbFCaBPs* (*Calflagin Tb17, Tb24*, and *Tb44*) in *T. brucei* did not affect parasite growth and motility (Emmer *et al*., 2010). The effects of *FCaBP* loss-of-function may differ between monoxenous parasites like *L. passim* (infecting only honey bees) and dixenous parasites like *T. brucei* (infecting both tsetse flies and mammals). It is also possible that the small amount of TbFCaBPs remaining after gene knockdown (Emmer *et al*., 2010) is sufficient to support the normal growth and motility of *T. brucei* in the culture medium. FCaBP likely functions as a calcium signaling protein, and it will be important to uncover how downstream effectors are involved in the growth and flagellar formation of *L. passim*.

### Roles of the flagellum in the host infection of *L. passim*

We observed that *LpFCaBPs*-deleted parasites can infect honey bee hosts as effectively as wild-type parasites in our assay (Fig. 7G). Although the mutant parasites are less active, they can reach the honey bee hindgut through normal gut flow. *L. passim* appears to attach to the hindgut wall through the structurally modified flagellum (Buendía-Abad *et al*., 2022), and the short flagella of the mutant parasites would be sufficient for attachment. In contrast, TbFCaBPs-depleted *T. brucei* showed attenuated parasitemia in infected mice (Emmer *et al*., 2010), but the underlying mechanism and the infection in the tsetse fly gut were not clarified. The precise *in vivo* roles of FCaBP in infecting insects and mammalian hosts remain to be answered.

### Establishing *L. passim* to express the GFP fusion proteins

To express the GFP fusion proteins LpFCaBP1-GFP, LpFCaBP2-GFP, and LpPFRP5-GFP, we amplified the complete ORFs of *LpFCaBP1*, *LpFCaBP2*, and *LpPFRP5* genes using PCR with KOD-FX DNA polymerase (TOYOBO), *L. passim* genomic DNA, and the primer pairs: LpFCaBP1-5 and LpFCaBP1-3, LpFCaBP2-5 and LpFCaBP2-3, and LpPFRP5-5 and LpPFRP5-3. The PCR products were digested with XbaI, purified from the gel, and then cloned into the XbaI site of pTrex-n-eGFP plasmid (Peng et al., 2014). To construct plasmids expressing LpFCaBP1 and LpFCaBP2 chimeric GFP fusion proteins, we amplified DNA fragments encoding the N-terminal 16 or 28 amino acids of LpFCaBP1 and LpFCaBP2 using 1444F primer and FCaBP1-N16-Rev, FCaBP1-N28-Rev, FCaBP2-N16-Rev, or FCaBP2-N28-Rev primer. Similarly, we amplified DNA fragments encoding amino acids 17-154 or 29-154 of LpFCaBP1 and LpFCaBP2 using FCaBP-487R primer and FCaBP1-N17-For, FCaBP1-N29-For, FCaBP2-N17-For, or FCaBP2-N29-For primer. To make the four chimeras, two DNA fragments for the corresponding N-terminal and internal amino acids of FCaBP1 and 2 or vice versa were ligated by fusion PCR using 1444F and FCaBP-487R primers. The resulting fusion PCR products and plasmids expressing LpFCaBP1-GFP or LpFCaBP2-GFP were digested with BamHI and ligated. Plasmid DNAs expressing GFPs with N-terminal sorting sequences of LpFCaBP1, LpFCaBP2, TcFCaBP, TbFCaBP, LdFCaBP1, CfFCaBP1, CfFCaBP2, or LsFCaBP1 were constructed by cloning the corresponding complementary oligonucleotides into the XbaI site of pTrex-n-eGFP.

We collected actively growing *L. passim* (4 × 10^7^), washed twice with 5 mL PBS, and then resuspended in 0.4 mL of Cytomix buffer without EDTA (20 mM KCl, 0.15 mM CaCl^2^, 10 mM K_2_HPO_4_, 25 mM HEPES and 5 mM MgCl_2_, pH 7.6) (Van den Hoff, Moorman and Lamers, 1992; Ngo *et al*., 1998). The parasites were electroporated twice (1 min interval) with 10 μg of each plasmid DNA constructed above using a Gene Pulser X cell electroporator (Bio-Rad) and cuvette (2 mm gap). We set the voltage, capacitance, and resistance at 1.5 kV, 25 μF, and infinity, respectively. The electroporated parasites were cultured in 4 mL of modified FP-FB medium (Salathé *et al*., 2012), and then G418 (200 μg/mL, Sigma-Aldrich) was added after 24 h to select the G418-resistant clones. To take the photos, we washed live *L. passim* expressing the GFP fusion protein three times with 1 mL PBS, and then mounted them on poly-L-lysine coated slide glass.

### Analysis of microsynteny with FCaBP genes in trypanosomatid parasites

We used TBLASTN and LpFCaBP1 as a query to identify the genomic contigs containing *FCaBP* in trypanosomatid parasites. The order and orientation of genes in the microsynteny with *FCaBP* of five *Leishmania* species, *L. seymouri*, and *L. pyrrhocoris* were determined based on annotated gene lists. For *L. passim, C. fasciculata, C. bombi*, and *C. expoeki*, we performed the analysis using TBLASTN. The amino acid sequences of 11 FCaBPs (Supplementary file 1) were aligned using Clustal Omega (https://www.ebi.ac.uk/Tools/msa/clustalo/).

### Testing interaction of N-terminal sorting sequence of LpFCaBP1 with either LpBBS1 or LpTULP

We constructed plasmid DNA for expressing ultraID with the N-terminal sorting sequence of LpFCaBP1. The DNA fragment encoding the sorting sequence was PCR amplified using 1444F and Ultra-Rev-FCaBP1 primers. We also PCR amplified the DNA fragment encoding ultraID using Ultra-For-FCaBP1 and Ultra-3-XhoI primers and used pSF3-ultraID (Kubitz et al., 2022) (ADDGENE: #172878) as a template. The two DNA fragments were ligated through fusion PCR followed by BamHI and XhoI digestion and cloned into the pTrex-n-eGFP plasmid at the same restriction enzyme sites. We first constructed plasmid DNA carrying triple Flag tags by inserting the annealed complementary oligonucleotides (3Flag-5 and 3Flag-3) into the XbaI and HindIII sites of tdTomato/pTREX-b (Lander et al., 2015) (ADDGENE: #68709). Then, we PCR amplified the entire ORFs of LpBBS1 and LpTULP (Supplementary file 2) using BBS1-5-HindIII and BBS1-3-ClaI stop primers as well as TULP-5-HindIII and TULP-3-ClaI stop primers. After digesting with HindIII and ClaI, the PCR products were cloned in the same sites of the above plasmid DNA with Flag tags.

*L. passim* was electroporated as described above with 10 μg each of LpFCaBP1N16-ultraID and either Flag-LpBBS1 or Flag-LpTULP. The parasites expressing both proteins were selected using G418 and blasticidin (50 μg/mL, Macklin). We detected the expressed proteins by immunofluorescence. The parasites were washed and mounted on a poly-L-lysine-coated 8-well chamber slide as above, fixed with 4% paraformaldehyde, permeabilized with PBS containing 0.1 % TX-100 (PT), and blocked with PT containing 5 % normal goat serum (PTG). The samples were incubated overnight at 4 ℃ with rabbit anti-Myc (500-fold dilution for LpFCaBP1N16-ultraID) or rabbit anti-Flag epitope (500-fold dilution for Flag-LpBBS1 and Flag-LpTULP) polyclonal antibodies (Proteintech) in PTG. The samples were washed five times with PT (5 minutes for each wash) and then incubated with Alexa Fluor 488 (for LpFCaBP1N16-ultraID) or Alexa Fluor 555 (for Flag-LpBBS1 and Flag-LpTULP) anti-rabbit IgG antibody (ThermoFisher) for 2 hours at room temperature. After another round of washing, the samples were observed under a microscope.

We incubated the parasites (10^8^) in 10 mL culture medium with or without 50 μM biotin for 3 hours at 28 ℃, followed by washing with 5 mL PBS twice. The samples were then suspended in 2 mL RIPA buffer (20 mM Tris-HCl, pH 7.5, 150 mM NaCl, 1 % NP-40, 0.5 % sodium deoxycholate, 0.1 % SDS) containing a protease inhibitor cocktail (Beyotime) and sonicated three times (5 seconds each) at amplitude 10 using a Q700 sonicator (Qsonica). After centrifugation, the supernatants were collected, and 50 μL of streptavidin agarose (Yeasen) and 12.5 μL of DYKDDDDK (Flag) Fab-Trap Agarose (Chromotek) were added to the samples with and without biotin treatment, respectively. The beads were washed five times (5 minutes each) with washing buffer (10 mM Tris-HCl, pH 7.5, 150 mM NaCl, 0.05 % NP-40, 0.5 mM EDTA) and then suspended in 70 μL of SDS-PAGE sample buffer (2 % SDS, 10 % glycerol, 10 % β-mercaptoethanol, 0.25 % bromophenol blue, 50 mM Tris-HCl, pH 6.8). The samples were heated at 95 ℃ for 5 minutes, and then 35 μL of each sample was applied to a 10 % SDS-PAGE gel. The proteins were transferred to a nitrocellulose membrane (Pall® Life Sciences) and the membrane was blocked with PBST (PBS with 0.1 % Tween-20) containing 5 % skim milk at room temperature for 30 minutes. The membrane was incubated with a 1000-fold diluted anti-Flag tag antibody (Proteintech) overnight at 4 ℃. After washing five times with PBST (5 minutes each), the membrane was incubated with a 10,000-fold diluted IRDye® 680RD donkey anti-rabbit IgG (H+L) secondary antibody (LI-COR Biosciences) in PBST containing 5 % skim milk at room temperature for 2 hours. Following another round of washing, the membrane was visualized using ChemiDoc MP (BioRad).

### Deletion of *LpFCaBP* genes by CRISPR

To delete *LpFCaBP1* and *2* genes, we designed the gRNA sequence using a custom gRNA design tool (http://grna.ctegd.uga.edu) (Peng and Tarleton, 2015). Complementary oligonucleotides (0.1 nmole each) corresponding to the sgRNA sequences (LpFCaBP-For and LpFCaBP-Rev) were phosphorylated by T4 polynucleotide kinase (TAKARA), annealed, and cloned into BbsI-digested pSPneogRNAH vector (Zhang and Matlashewski, 2015). We electroporated *L. passim* expressing Cas9 (Liu, Lei and Kadowaki, 2019) with 10 μg of the constructed plasmid DNA and selected the transformants by blasticidin and G418 to establish parasites expressing both Cas9 and *LpFCaBP* gRNA.

For the construction of donor DNA for *LpFCaBP* genes, we performed fusion PCR of three DNA fragments: 5’UTR of *LpFCaBP1* (574 bp, LpFCaBP1 5’UTR-F and LpFCaBP1 5’UTR-R), the ORF of the *Hygromycin B phosphotransferase (Hph)* gene derived from pCsV1300 (Park et al., 2013) (1026 bp, LpFCaBP1 Hph-F and LpFCaBP2 Hph-R), and 3’UTR of *FCaBP2* (590 bp, LpFCaBP2 3’UTR-F and LpFCaBP2 3’UTR-R). The fusion PCR products were cloned into the EcoRV site of pBluescript II SK(+) and the linearized plasmid DNA (10 μg) by HindIII was used for electroporation of *L. passim* expressing both Cas9 and *LpFCaBP* gRNA as described above.

After electroporation, *L. passim* resistant to blasticidin, G418, and hygromycin (150 μg/mL, Sigma-Aldrich) were selected, and single parasites were cloned by serial dilutions in a 96-well plate. The genotype of each clone was initially determined through the detection of 5’ wild-type (WT) and knock-out (KO) alleles for *LpFCaBPs* by PCR. After identifying heterozygous (+/-) and homozygous (-/-) KO clones, and their 5’WT (LpFCaBP1 5’UTR-Outer-F and LpFCaBP-152R), 5’KO (LpFCaBP1 5’UTR-Outer-F and Hyg-159R), 3’WT (LpFCaBP-538F and LpFCaBP2 3’UTR-Outer-R), and 3’KO (Hyg-846F and LpFCaBP2 3’UTR-Outer-R) alleles were confirmed by PCR using the specific primer sets.

### RT-PCR

Total RNA was extracted from wild-type, *LpFCaBP*s heterozygous, and homozygous mutant parasites using TRIzol reagent (Sigma-Aldrich). Reverse transcription of 0.2 μg of total RNA was performed using ReverTra Ace (TOYOBO) and random primers, followed by PCR with KOD-FX DNA polymerase. To specifically detect *FCaBP1* and *2* mRNAs, we designed reverse primers corresponding to the N-terminal sorting sequences (FCaBP1-36R and FCaBP2-36R). *L. passim* splice leader sequence (LpSL-F) was used as the forward primer. To detect *LpGAPDH* mRNA, we used LpGAPDH-F and LpGAPDH-R primers.

### Culture, Flagellar length measurement, and motility measurement of *L. passim*

For culture, flagellar length measurement, and motility measurement, we inoculated wild-type and *LpFCaBPs* homozygous mutant parasites in the culture medium at 10^4^/mL. The number of parasites was counted every 12 hours using a hemocytometer, and images of the cultured parasites were captured simultaneously for a week. The length of the flagellum of individual parasites was measured at 60 hours after culture using Image-J. To record the movement of parasites, videos were created by taking images for 1 minute every second. The movement of parasites was tracked using TrackMate v7.10.2 (Ershov et al., 2022) as a Fiji (Schindelin et al., 2012) plugin. The videos were imported to Fiji, converted to 8-bit grayscale, and the brightness and contrast were adjusted for better tracking. The Laplacian of Gaussian detector was used with an estimated object diameter of 30.0-36.0 pixels and a quality threshold of 0.2-0.5 for the detection of individual parasites. We used the Simple Linear Assignment Problem tracker to track the parasites by adjusting the linking maximum distance and gap-closing maximum distance to 35.0-200.0 pixels and gap-closing maximum frame gap to 1.

### Honey bee infection

To infect honey bees with *L. passim*, parasites collected during the logarithmic growth phase (5 × 10^5^/mL) were washed with PBS and suspended in sterile 10 % sucrose/PBS at 5 × 10^4^/μL. Newly emerged honey bee workers were collected by placing the frames with late pupae in 33 °C incubator and starved for 2-3 hours. Twenty individual honey bees were fed with 2 μL of the sucrose/PBS solution containing either wild-type or *LpFCaBPs* homozygous mutant (10^5^ parasites in total). The infected honey bees were maintained in metal cages at 33 °C for 14 days and then frozen at -80 °C. This experiment was repeated three times. We sampled eight honey bees from each of the above three experiments, and thus analyzed 24 honey bees in total infected with either wild-type or *LpFCaBPs* homozygous mutant. Genomic DNAs were extracted from the whole abdomens of individual bees using DNAzol® reagent (Thermo Fisher). We quantified *L. passim* in the infected honey bee by qPCR using LpITS2-F and LpITS2-R primers which correspond to the part of internal transcript spacer region 2 (ITS2) in the *ribosomal RNA* gene. Honey bee *AmHsTRPA* was used as the internal reference using AmHsTRPA-F and AmHsTRPA-R primers (Liu *et al*., 2020). The relative abundances of *L. passim* in the individual honey bees (24 each infected by wild-type or mutant *L. passim*) were calculated by the ΔC_t_ method and we set one sample infected by wild-type as 1. The statistical analysis was performed using the Brunner-Munzel test. All of the above primers are listed in Supplementary file 3.

## Acknowledgements

We thank Yizhen Shao for his contribution to this study.

## Funding

This work was supported by Jinji Lake Double Hundred Talents Programme to TK.

## Author contribution

TK conceived and designed research strategy and wrote the paper. XY performed the experiments. XY and TK analyzed data.

---

**Supplementary file 1 DNA and amino acid sequences of 18 FCaBPs as well as the aligned sequences**

**Supplementary file 2 DNA and amino acid sequences of three *Trypanosoma theileri***

**FCaBPs, LpPFRP5, LpBBS1, and LpTULP**

**Supplementary file 3 List of primers used in this study Supplementary video 1 Movement of wild-type *L. passim***

**Supplementary video 2 Movemnt of FCaBPs deleted mutant *L. passim***

## Notes

### Competing Interest Statement

The authors have declared no competing interest.

