## Supplementary file 1 for "Protein subcellular relocalization and function of duplicated flagellar calcium binding protein genes in honey bee trypanosomatid parasite"

**Alignment of eight FCaBPs**

TcFCaBP	**MGACGSKGS----TSDKGLASDKDG** **KNA**KDRKEAWERIRQAIPREKTAEAKQRRIELFKK	56

TbFCaBP	**MG-CSGSKNTTNSKDGAASKGGKDG** KTTADRKVAWERIRCAIPRDKDAESKSRRIELFKQ	59

LdFCaBP1	**MG-CNAT---------KAARKPSED** KTAADRKVAWEKICQRLPRQKTPEDKELHIELFKR	50

CfFCaBP1	**MG-CASSAF----SS----KSKKEG** KNASDRKAAWEGIRERLPRKKTDEDKERRIELFKK	51

CfFCaBP2	**MG-CISSKS----TQ----TGKKEG** KTAAERKAAWEGIRQRLPRRKTAEDKARRIELFKK	51

LsFCaBP1	**MG-CASSTS----SS---KGGKKEG** KSAAERRAVWGSVRQSLPRLKTAVDKERRIALFKE	52

LpFCaBP1	**MG-CASSLF----SS----KSKTED** KTAAERKVAWEKIRERLPRRKTPEDKERRIELFKK	51

LpFCaBP2	**MG-CISSKS----TQ----IGKKEC** KTAAERKAAWEGIRQRLPRRKTPEDKQRRIELFKK	51

** * .. .: *.: :*: .* : :** *	* :* ***.

TcFCaBP	FDKNETGKLCYDEVYSGCLEVLKLDEFTSRVRDITKRAFDKSRTLGSKLE-NKGSEDFVE	115

TbFCaBP	FDTNGTGKLGFREVLDGCYSVLKLDEFTTHLPDIVQRAFDKAKDLGNKVK-GVGEEDLVE	118

LdFCaBP	FCQSDTEKLTMEEVYQGCVSILQLDEFTTRLRSIVKRAFTKAKSMGNTSGVGQCDDEVVE	110

CfFCaBP1	FDQNNTGKLSFEETYKGCVEILHLDEFTTRLRDIVKRAFNKAKDMGTKDR-DAGSAQFVE	110

CfFCaBP2	FDKNNTGKLSMEETYEGCVSVLHLDEFTTRLRDIVKRAFNKAKDMGTKDR-DAGSAQFVE	110

LsFCaBP	FDKNNSGKLSMEEVYKGCVNVLHLDEFTTRLRDIVKRAFNKAKDMGTKER-NEGSSDFVE	111

LpFCaBP1	FDQNNTNRLSVEEVYEGCVDILHLDEFTTRLRDIVKRAFNKAKEMGTKDR-GAGSDKFVE	110

LpFCaBP2	FDKNNTNRLTLEEVYEGCVSILHLDEFTTRLRDIVKRAFNKAKEMGTKDR-GAGSDKFVE	110

* . : :*	*. .** .:*:*****::: .*.:*** *:: :*.. . . ..**

TcFCaBP	FLEFRLMLCYIYDFFELTVMFDEIDASGNMLVDEEEFKRAVPKLEAWGAKVEDPAALFKE	175

TbFCaBP	FLEFRLMLCYIYDIFELTVMFDTMDKDGSLLLELQEFKEALPKLKEWGVDITDATTVFNE	178

LdFCaBP	FLEFRLILCYIYDYLELTVMFDEIDTSGNMLVDAREFKAAVPKMGEWGLVIEDPDTIFKE	170

CfFCaBP1	FLEFRLMLCYIYDYFELTVMFDEIDTSGNMLIDAKEFKAAVPKIAEWGLKINDPDAVFKK	170

CfFCaBP2	FLEFRLMLCYIYDYFELTVMFDEIDTSGNMLIDAKEFKKAAPKITEWGVKITDADAVFKE	170

LsFCaBP	FLEFRLLLCYIYDYFELTVMFDEIDTSGNMLIDANEFQKAVPKIAEWGLEISDPDAVFKE	171

LpFCaBP1	FLEFRLMLCYIYDYFELTVMFDEIDTSGNMLIDAKEFKKAVPRIQEWGVKIEDPDAVFKE	170

LpFCaBP2	FLEFRLMLCYIYDYFELTVMFDEIDTSGNMLIDAKEFKKAVPRIQEWGVKIEDPDAVFKE	170

******:****** :******* :* .*.:*:: .**: * *:: ** : * ::*::

TcFCaBP	LDKNGTGSVTFDEFAAWASAVKLDADGDPDNVPESA----	211

TbFCaBP	IDTNGSGVVTFDEFSCWAVTKKLQVCGDPDGEENGANEGN	218

LdFCaBP	IDDNGSGQVPFDELAAWASRSSAGH---------------	195

CfFCaBP1	IDDNGSGQVTFDEFAAWAAAQKLDADENPGNTE-------	203

CfFCaBP2	IDDNGSGQVTFDEFAAWASARKLDADGDPDNT--------	202

LsFCaBP	IDDNGSGQITFDEFAVWAAAHKLDADGDPDNTAS------	205

LpFCaBP1	IDNNGSGQITFDEFAAWAAAHKLDADENPDNTE-------	203

LpFCaBP2	IDSNGSGQMTFDEFAAWASAHKLDADGDPDNMAA------	204

:* **:* : ***:: **	.

**Alignment of 11 FCaBPs**

TcFCaBP	MGACGSKGS----TSDKGLASDKDG KNAKDRKEAWERIRQAIPREKTAEAKQRRIELFKK	56

TbFCaBP	MG-CSGSKNTTNSKDGAASKGGKDG KTTADRKVAWERIRCAIPRDKDAESKSRRIELFKQ	59

LdFCaBP1	MG-CNATK---------AARKPSED KTAADRKVAWEKICQRLPRQKTPEDKELHIELFKR	50

CfFCaBP1	MG-CASSAF----SS----KSKKEG KNASDRKAAWEGIRERLPRKKTDEDKERRIELFKK	51

CfFCaBP2	MG-CISSKS----TQ----TGKKEG KTAAERKAAWEGIRQRLPRRKTAEDKARRIELFKK	51

LepFCaBP1	MG-CASSAF----SS----KSKKEG KTAAERKAAWESIRQRLPRQKTAEDKERRIELFKK	51

LepFCaBP2-A	MG-CTSSKS----TQ----TGKKEG KTAAERKAAWESIRQRLPRQKTAEDKERRIELFKK	51

LepFCaBP2-B	MG-CTSSKS----TQ----TGKKEG KTAAERKAAWESIRQRLPRQKTAEDKERRIELFKK	51

LsFCaBP1	MG-CASSTS----SS---KGGKKEG KSAAERRAVWGSVRQSLPRLKTAVDKERRIALFKE	52

LpFCaBP1	MG-CASSLF----SS----KSKTED KTAAERKVAWEKIRERLPRRKTPEDKERRIELFKK	51

LpFCaBP2	MG-CISSKS----TQ----IGKKEC KTAAERKAAWEGIRQRLPRRKTPEDKQRRIELFKK	51

** * .. .: *.: :*: .* : :** *	* :* ***.

TcFCaBP	FDKNETGKLCYDEVYSGCLEVLKLDEFTSRVRDITKRAFDKSRTLGSKLE-NKGSEDFVE	115

TbFCaBP	FDTNGTGKLGFREVLDGCYSVLKLDEFTTHLPDIVQRAFDKAKDLGNKVK-GVGEEDLVE	118

LdFCaBP1	FCQSDTEKLTMEEVYQGCVSILQLDEFTTRLRSIVKRAFTKAKSMGNTSGVGQCDDEVVE	110

CfFCaBP1	FDQNNTGKLSFEETYKGCVEILHLDEFTTRLRDIVKRAFNKAKDMGTKDR-DAGSAQFVE	110

CfFCaBP2	FDKNNTGKLSMEETYEGCVSVLHLDEFTTRLRDIVKRAFNKAKDMGTKDR-DAGSAQFVE	110

LepFCaBP1	FDQNGSGKLTFEETYKGCVSVLHLDEFTTRLRDIVKRAFNKAKDMGTKER-NSGSEQFVE	110

LepFCaBP2-A	FDQNGSGKLTFEETYKGCVSVLHLDEFTTRLRDIVKRAFNKAKDMGTKER-NSGSEQFVE	110

LepFCaBP2-B	FDQNGSGKLTFEETYKGCVSVLHLDEFTTRLRDIVKRAFNKAKDMGTKER-NSGSEQFVE	110

LsFCaBP1	FDKNNSGKLSMEEVYKGCVNVLHLDEFTTRLRDIVKRAFNKAKDMGTKER-NEGSSDFVE	111

LpFCaBP1	FDQNNTNRLSVEEVYEGCVDILHLDEFTTRLRDIVKRAFNKAKEMGTKDR-GAGSDKFVE	110

LpFCaBP2	FDKNNTNRLTLEEVYEGCVSILHLDEFTTRLRDIVKRAFNKAKEMGTKDR-GAGSDKFVE	110

* . : :*	*. .** .:*:*****::: .*.:*** *:: :*.. . . ..**

TcFCaBP	FLEFRLMLCYIYDFFELTVMFDEIDASGNMLVDEEEFKRAVPKLEAWGAKVEDPAALFKE	175

TbFCaBP	FLEFRLMLCYIYDIFELTVMFDTMDKDGSLLLELQEFKEALPKLKEWGVDITDATTVFNE	178

LdFCaBP1	FLEFRLILCYIYDYLELTVMFDEIDTSGNMLVDAREFKAAVPKMGEWGLVIEDPDTIFKE	170

CfFCaBP1	FLEFRLMLCYIYDYFELTVMFDEIDTSGNMLIDAKEFKAAVPKIAEWGLKINDPDAVFKK	170

CfFCaBP2	FLEFRLMLCYIYDYFELTVMFDEIDTSGNMLIDAKEFKKAAPKITEWGVKITDADAVFKE	170

LepFCaBP1	FLEFRLMLCYIYDYFELTIMFDEIDTSGNMLIDAKEFKKAAPKIAEWGVKISDPDAVFKE	170

LepFCaBP2-A	FLEFRLMLCYIYDYFELTIMFDEIDTSGNMLIDAKEFKKAAPKIAEWGVKISDPDAVFKE	170

LepFCaBP2-B	FLEFRLMLCYIYDYFELTIMFDEIDTSGNMLIDAKEFKKAAPKIAEWGVKISDPDAVFKE	170

LsFCaBP1	FLEFRLLLCYIYDYFELTVMFDEIDTSGNMLIDANEFQKAVPKIAEWGLEISDPDAVFKE	171

LpFCaBP1	FLEFRLMLCYIYDYFELTVMFDEIDTSGNMLIDAKEFKKAVPRIQEWGVKIEDPDAVFKE	170

LpFCaBP2	FLEFRLMLCYIYDYFELTVMFDEIDTSGNMLIDAKEFKKAVPRIQEWGVKIEDPDAVFKE	170

******:****** :***:*** :* .*.:*:: .**: * *:: ** : * ::*::

TcFCaBP	LDKNGTGSVTFDEFAAWASAVKLDADGDPDNVPESA----	211

TbFCaBP	IDTNGSGVVTFDEFSCWAVTKKLQVCGDPDGEENGANEGN	218

LdFCaBP1	IDDNGSGQVPFDELAAWASRSSAGH---------------	195

CfFCaBP1	IDDNGSGQVTFDEFAAWAAAQKLDADENPGNTE-------	203

CfFCaBP2	IDDNGSGQVTFDEFAAWASARKLDADGDPDNT--------	202

LepFCaBP1	IDDNGSGQITFDEFAAWASAHKLDVDGDPDNAE-------	203

LepFCaBP2-A	IDDNGSGQITFDEFAAWASAHKLDVDGDPDNVAA------	204

LepFCaBP2-B	IDDNGSGQITFDEFAAWASAHKLDVDGDPDNAE-------	203

LsFCaBP1	IDDNGSGQITFDEFAVWAAAHKLDADGDPDNTAS------	205

LpFCaBP1	IDNNGSGQITFDEFAAWAAAHKLDADENPDNTE-------	203

LpFCaBP2	IDSNGSGQMTFDEFAAWASAHKLDADGDPDNMAA------	204

:* **:* : ***:: **	.

>TcFCaBP

MGACGSKGSTSDKGLASDKDGKNAKDRKEAWERIRQAIPREKTAEAKQRRIELFKKFDKNETGKLCYDEV

YSGCLEVLKLDEFTSRVRDITKRAFDKSRTLGSKLENKGSEDFVEFLEFRLMLCYIYDFFELTVMFDEID

ASGNMLVDEEEFKRAVPKLEAWGAKVEDPAALFKELDKNGTGSVTFDEFAAWASAVKLDADGDPDNVPES

A

ATGGGTGCTTGTGGGTCGAAGGGCTCGACGAGCGACAAGGGGTTGGCGAGCGATAAGGACGGCAAGAACGCCA

AGGACCGCAAGGAAGCGTGGGAGCGCATTCGCCAGGCGATTCCTCGTGAGAAGACCGCCGAGGCAAAACA

GCGCCGCATCGAGCTCTTCAAGAAGTTCGACAAGAACGAGACCGGGAAGCTGTGCTACGATGAGGTGTAC

AGCGGCTGCCTCGAGGTGCTGAAGTTGGACGAGTTCACGTCGCGAGTGCGCGACATCACGAAGCGTGCAT

TCGACAAGTCGAGGACCCTGGGCAGCAAGCTGGAGAACAAGGGCTCCGAGGACTTTGTTGAATTTCTGGA

GTTCCGTCTGATGCTGTGCTACATCTACGACTTCTTCGAGCTGACGGTGATGTTCGACGAGATTGACGCC

TCCGGCAACATGCTGGTCGACGAGGAGGAGTTCAAGCGCGCCGTGCCCAAGCTTGAGGCGTGGGGCGCCA

AGGTCGAGGATCCCGCGGCGCTGTTCAAGGAGCTCGATAAGAACGGCACTGGGTCCGTGACGTTCGACGA

GTTTGCTGCGTGGGCTTCTGCAGTCAAACTGGACGCCGACGGCGACCCGGACAACGTGCCGGAGAGCGCG

TGA

>TbFCaBP

MGCSGSKNTTNSKDGAASKGGKDGKTTADRKVAWERIRCAIPRDKDAESKSRRIELFKQFDTNGTGKLGF

REVLDGCYSVLKLDEFTTHLPDIVQRAFDKAKDLGNKVKGVGEEDLVEFLEFRLMLCYIYDIFELTVMFD

TMDKDGSLLLELQEFKEALPKLKEWGVDITDATTVFNEIDTNGSGVVTFDEFSCWAVTKKLQVCGDPDGE

ENGANEGN

ATGGGTTGCTCAGGATCGAAGAATACAACGAACTCCAAGGATGGTGCCGCCAGTAAGGGTGGAAAGGACG

GGAAAACTACTGCGGACCGGAAGGTTGCGTGGGAGCGTATTCGGTGTGCGATTCCCCGTGACAAGGATGC

TGAGTCGAAAAGCCGTCGCATTGAACTCTTCAAGCAGTTTGACACGAACGGGACTGGAAAACTCGGCTTC

AGGGAGGTGCTTGATGGTTGCTACAGTGTACTCAAGTTGGATGAGTTCACGACTCACCTTCCTGACATTG

TCCAGCGTGCGTTTGACAAAGCGAAGGACCTCGGCAACAAGGTAAAGGGTGTGGGTGAAGAGGATCTTGT

GGAGTTCCTTGAATTCCGCCTGATGCTTTGCTACATCTATGATATCTTTGAGTTGACTGTAATGTTCGAC

ACCATGGATAAGGATGGAAGTTTGTTGCTTGAGCTACAGGAGTTCAAGGAAGCCTTGCCGAAATTGAAGG

AGTGGGGTGTTGATATCACTGACGCTACCACTGTGTTTAATGAGATCGACACTAATGGATCTGGTGTTGT

GACATTCGATGAATTCTCATGCTGGGCTGTCACTAAAAAGTTGCAGGTATGTGGCGATCCTGATGGTGAG

GAGAACGGAGCAAATGAAGGTAATTGA

>LiFCaBP1

MGCNATKAARKPSEDKTAAERKVAWEKICQRLPRQKTPEDKELRIELFKRFCQSDTEKLTMEEVYQGCVSILQLDEFTTRLRSIVKRAFTKAKSMGNTAGVGQCDDEFVEFLEFRLMLCYIYDYLELTVMFEEIDTSGNMLVDAREFKAAVPKMGEWGLVIEDPDTIFKEIDDNGSGQVPFDELAAWATARKLDADEQANNSE

ATGGGCTGCAACGCTACCAAGGCTGCCCGGAAGCCCAGCGAGGACAAGACTGCCGCTGAGCGCAAGGTCG

CGTGGGAGAAAATTTGTCAGCGCCTGCCCCGCCAGAAGACGCCCGAGGACAAGGAGCTCCGCATCGAGCT

TTTTAAGAGGTTCTGCCAGAGCGACACGGAAAAGCTCACGATGGAGGAGGTCTACCAGGGCTGTGTGAGC

ATCCTCCAGCTCGATGAGTTTACAACGCGCCTGCGCAGCATCGTGAAACGTGCCTTCACTAAGGCAAAGA

GCATGGGCAACACCGCCGGCGTCGGCCAATGCGACGACGAGTTTGTCGAATTTCTTGAGTTTCGGCTGAT

GCTGTGCTACATATACGACTACCTCGAGCTCACCGTCATGTTTGAAGAGATCGACACCTCCGGCAACATG

CTGGTGGACGCGAGGGAGTTCAAGGCCGCCGTGCCGAAGATGGGTGAGTGGGGCCTTGTCATTGAGGATC

CGGACACAATCTTCAAGGAAATCGACGACAACGGCTCCGGCCAGGTGCCCTTTGACGAGCTCGCTGCGTG

GGCCACTGCCCGCAAGCTTGATGCTGACGAGCAGGCTAACAACTCTGAGTAA

>LbFCaBP1

MGCNTSKAAPKSGEDRTAAERKAAWEKIRQSLPRQRTPEDKERHVLLFKKFDNSESGKLTMEEVYQGCVD

ILQLDEFTTRLRDIVQRAFSKAKSMGNTADDGQESSDYVELLEFRLMLCYIYDYFELTVMFDEIDTSGNM

LVSAKEFKAALPRIGEWGVAIEDPDKIFKEIDTNSTGQVTFDEFAAWATGCKLQSDADAMKSE

ATGGGCTGCAACACTTCCAAGGCTGCTCCGAAGTCTGGAGAGGATAGGACTGCCGCCGAGCGCAAGGCTG

CGTGGGAGAAGATCCGCCAGAGCCTACCCCGCCAGAGGACCCCTGAGGACAAGGAGCGCCACGTATTGTT

GTTCAAGAAGTTCGATAATAGCGAATCTGGGAAGCTCACAATGGAGGAGGTCTACCAGGGCTGCGTGGAT

ATTCTTCAGCTTGATGAGTTCACAACGCGCCTGCGCGACATCGTGCAGAGAGCCTTCAGCAAGGCCAAAA

GCATGGGCAACACCGCTGACGACGGCCAGGAAAGCTCTGACTACGTTGAGCTCTTGGAGTTCCGCCTGAT

GCTGTGCTACATCTACGACTACTTCGAGCTCACCGTCATGTTCGACGAGATTGACACCTCCGGCAATATG

CTGGTGAGCGCGAAAGAGTTCAAGGCTGCCCTGCCGAGAATTGGTGAGTGGGGCGTTGCCATCGAGGATC

CAGACAAAATATTCAAGGAGATCGACACGAACAGCACTGGGCAGGTGACATTTGACGAATTCGCTGCGTG

GGCCACTGGATGCAAGCTTCAATCTGACGCAGATGCTATGAAGTCCGAGTAG

>LmFCaBP1

MGCNATKAAGKSGEGKTAAERKVAWEKICQNLPRQRTPEDKERRSDLFKKFSQNDTEKLTMEEVYQGCVRILQLDEFTTRLHDIVKRAFNKAKSMANTAGDCQGDDEFVEFLEFRLMLCYIYDYFKLTVMFDEIDTSGNMLVDAKELKAAVPKIGEWGLVIEDPDTVFKQIDDNGSGQVSFNEFASWATARKLDADEEANDSE

ATGGGCTGTAACGCTACCAAGGCTGCCGGCAAGTCCGGCGAGGGCAAGACCGCCGCTGAGCGCAAGGTTG

CGTGGGAGAAAATTTGTCAGAACCTGCCTCGCCAGAGGACGCCTGAGGACAAGGAACGCCGCAGCGATCT

TTTCAAGAAGTTTTCCCAGAACGACACGGAAAAGCTCACGATGGAGGAAGTCTATCAAGGCTGTGTACGT

ATCCTCCAGCTCGATGAGTTCACGACGCGCCTGCACGACATTGTGAAGCGTGCCTTCAACAAGGCAAAGA

GCATGGCGAACACCGCCGGCGACTGCCAGGGCGACGACGAGTTTGTCGAATTTCTTGAGTTTCGGCTGAT

GCTGTGCTACATATACGACTACTTCAAGCTCACCGTTATGTTTGACGAGATCGACACCTCCGGCAACATG

CTGGTGGACGCGAAGGAGCTCAAGGCCGCCGTGCCGAAGATCGGTGAGTGGGGCCTTGTCATTGAGGATC

CAGACACAGTCTTCAAGCAAATCGACGACAACGGCTCCGGCCAGGTGTCCTTTAACGAGTTCGCTTCGTG

GGCGACTGCCCGCAAGCTTGATGCTGACGAGGAGGCTAACGACTCTGAGTAG

>LdFCaBP1

MGCNATKAARKPSEDKTAADRKVAWEKICQRLPRQKTPEDKELHIELFKRFCQSDTEKLTMEEVYQGCVS

ILQLDEFTTRLRSIVKRAFTKAKSMGNTSGVGQCDDEVVEFLEFRLILCYIYDYLELTVMFDEIDTSGNM

LVDAREFKAAVPKMGEWGLVIEDPDTIFKEIDDNGSGQVPFDELAAWASRSSAGH

ATGGGCTGCAACGCTACCAAGGCTGCCCGGAAGCCCAGCGAGGACAAGACTGCCGCTGAGCGCAAGGTCG

CGTGGGAGAAAATTTGTCAGCGCCTGCCCCGCCAGAAGACGCCCGAGGACAAGGAGCTCCGCATCGAGCT

TTTTAAGAGGTTCTGCCAGAGCGACACGGAAAAGCTCACGATGGAGGAGGTCTACCAGGGCTGTGTGAGC

ATCCTCCAGCTCGAGGAGTTTACAACGCGCCTGCGCAGCATCGTGAAACGTGCCTTCACTAAGGCAAAGA

GCATGGGCAACACCGCCGGCGTCGGCCAATGCGACGACGAGTTTGTCGAATTTCTTGAGTTTCGGCTGAT

GCTGTGCTACATATACGACTACCTCGAGCTCACCGTCATGTTTGACGAGATCGACACCTCCGGCAACATG

CTGGTGGACGCGAGGGAGTTCAAGGCCGCCGTGCCGAAGATGGGTGAGTGGGGCCTTGTCATTGAGGATC

CGGACACAATCTTCAAGGAAATCGACGACAACGGCTCCGGCCAGGTGTCCTTTGACGAGCTCGCTGCGTG

GGCCACTGCCCGCAAGCTTGATGCTGACGAGCAGGCTAACAACTCTGAGTAA

>CfFCaBP1

MGCASSAFSSKSKKEGKNASDRKAAWEGIRERLPRKKTDEDKERRIELFKKFDQNNTGKLSFEETYKGCVEILHLDEFTTRLRDIVKRAFNKAKDMGTKDRDAGSAQFVEFLEFRLMLCYIYDYFELTVMFDEIDTSGNMLIDAKEFKAAVPKIAEWGLKINDPDAVFKKIDDNGSGQVTFDEFAAWAAAQKLDADENPGNTE

ATGGGTTGCGCTTCTTCCGCTTTTTCCTCCAAGTCCAAGAAGGAGGGCAAGAACGCCAGCGATCGCAAGGCCGCGTGGGAGGGCATTCGCGAGCGTCTGCCGCGCAAGAAGACTGACGAGGACAAGGAGCGCCGCATTGAGCTCTTCAAGAAGTTCGACCAGAACAACACCGGCAAGCTGTCGTTTGAGGAGACGTACAAGGGATGCGTAGAGATTCTGCACCTGGACGAGTTCACCACTCGTCTGCGCGACATCGTCAAGCGCGCCTTCAACAAGGCCAAGGACATGGGCACGAAGGACCGCGATGCGGGCAGTGCTCAGTTCGTGGAGTTCCTCGAGTTCCGTCTGATGCTCTGCTACATCTACGACTACTTTGAGCTGACCGTGATGTTTGACGAGATCGACACTTCTGGCAACATGCTGATCGACGCGAAGGAGTTCAAGGCCGCCGTCCCTAAGATTGCGGAGTGGGGATTGAAGATTAATGACCCTGATGCCGTCTTCAAGAAGATCGACGACAACGGCTCCGGTCAGGTGACCTTCGACGAGTTTGCAGCGTGGGCGGCGGCCCAGAAACTCGACGCTGACGAGAACCCTGGCAACACGGAATAA

>CfFCaBP2

MGCISSKSTQTGKKEGKTAAERKAAWEGIRQRLPRRKTAEDKARRIELFKKFDKNNTGKLSMEETYEGCVSVLHLDEFTTRLRDIVKRAFNKAKDMGTKDRDAGSAQFVEFLEFRLMLCYIYDYFELTVMFDEIDTSGNMLIDAKEFKKAAPKITEWGVKITDADAVFKEIDDNGSGQVTFDEFAAWASARKLDADGDPDNTAA

ATGGGGTGCATTTCTTCCAAGTCTACTCAGACCGGCAAGAAGGAGGGCAAGACTGCCGCCGAGCGCAAGGCCGCGTGGGAGGGCATTCGCCAGCGCCTGCCGCGCCGCAAGACTGCCGAGGACAAGGCACGCCGCATTGAGCTCTTCAAGAAGTTCGATAAGAACAACACCGGCAAGCTCAGCATGGAGGAGACGTACGAGGGCTGCGTGTCTGTGTTGCACCTGGACGAGTTCACCACTCGTCTGCGCGACATCGTCAAGCGCGCCTTCAACAAGGCCAAGGACATGGGCACGAAGGACCGCGATGCGGGCAGCGCTCAGTTCGTGGAGTTCCTCGAGTTCCGTCTGATGCTCTGCTACATCTACGACTACTTCGAGCTGACCGTGATGTTTGACGAGATCGACACTTCTGGCAACATGCTGATCGACGCGAAGGAGTTCAAGAAGGCTGCTCCCAAGATTACGGAGTGGGGCGTGAAGATCACCGATGCCGATGCCGTCTTCAAGGAGATCGACGACAACGGCTCCGGTCAGGTGACCTTCGACGAGTTTGCGGCGTGGGCGAGCGCGCGCAAGCTCGACGCCGACGGCGATCCGGATAACACCGCTGCTTAA

>LsFCaBP1

MGCASSTSSSKGGKKEGKSAAERRAVWGSVRQSLPRLKTAVDKERRIALFKEFDKNNSGKLSMEEVYKGC

VNVLHLDEFTTRLRDIVKRAFNKAKDMGTKERNEGSSDFVEFLEFRLLLCYIYDYFELTVMFDEIDTSGN

MLIDANEFQKAVPKIAEWGLEISDPDAVFKEIDDNGSGQITFDEFAVWAAAHKLDADGDPDNTAS

ATGGGTTGCGCCTCTTCCACATCCTCTTCAAAGGGCGGCAAGAAGGAGGGCAAGAGCGCCGCAGAGCGCA

GAGCTGTGTGGGGCAGCGTTCGCCAGAGTCTGCCGCGCCTGAAGACTGCAGTGGATAAGGAGCGCCGTAT

TGCACTCTTTAAGGAGTTCGATAAGAACAACAGTGGTAAGCTCAGCATGGAGGAGGTGTACAAGGGTTGC

GTGAATGTGCTGCATCTGGATGAGTTCACGACTCGCCTGCGCGACATCGTGAAGCGCGCCTTCAACAAGG

CCAAGGATATGGGAACGAAGGAGCGCAACGAGGGCAGCTCGGACTTCGTGGAGTTCCTGGAGTTCCGTCT

GTTGCTGTGCTACATCTACGACTACTTTGAGCTGACGGTGATGTTCGATGAGATCGACACTTCTGGCAAC

ATGCTGATCGATGCGAATGAGTTCCAGAAGGCTGTCCCGAAGATCGCGGAGTGGGGTTTGGAGATTTCCG

ACCCCGATGCCGTGTTCAAGGAGATCGACGACAACGGCTCTGGTCAGATCACGTTTGACGAGTTCGCCGT

ATGGGCTGCCGCCCACAAGCTGGATGCTGATGGCGATCCGGATAATACCGCTTCGTAA

>CeFaBP1

MGCASSVFSSKSKKEGKSAAERKVAWGRIRERLPRKKTPEDKERRIELFKKFDQNGTGKLSFEETYRGCVDILHLDEFTTRLRDIVKRAFSKAKDMGTKDRGAGSDQFVEFLEFRLMLCYIYDYFELTVMFDEIDTSGNMLIDAKEFKAAVPKIAEWGLKIEDPDAVFKQIDDNGSGQITFDEFAAWAAARKLDADENPNNAE

ATGGGCTGCGCCTCCTCCGTATTTTCCTCCAAGTCAAAGAAAGAGGGCAAGAGCGCCGCCGAACGTAAAGTGGCGTGGGGGAGGATCCGCGAGCGTCTGCCGCGCAAGAAGACTCCTGAGGACAAAGAGCGCCGCATTGAACTCTTCAAGAAGTTCGATCAGAACGGCACCGGTAAACTTTCGTTTGAGGAGACGTACAGGGGGTGCGTAGACATTCTGCATCTGGATGAGTTCACGACTCGCCTGCGCGACATCGTGAAGCGCGCCTTCAGCAAGGCCAAAGACATGGGCACGAAGGACCGCGGCGCGGGTAGCGACCAGTTTGTGGAGTTCCTTGAGTTCCGCCTGATGCTGTGCTACATCTACGACTACTTTGAGCTGACTGTGATGTTCGACGAGATCGATACTTCTGGTAACATGCTTATCGACGCGAAGGAGTTCAAGGCCGCGGTCCCCAAGATTGCGGAGTGGGGCCTCAAGATTGAAGATCCCGACGCCGTGTTCAAGCAGATCGATGACAACGGCTCTGGCCAGATCACGTTCGACGAGTTCGCTGCGTGGGCAGCTGCGCGCAAGCTCGATGCTGACGAGAACCCCAACAACGCTGAGTGA

>CeFCaBP2

MGCISSKSTSSKKEGKNAAERKVAWEGIRQRLPRRKTPEDKARRIELFKQFDKNNTGKLSLEEVYNGCVTVLHLDEFTTRLRDIVKRAFSKAKDMGTKDRGAGSDQFVEFLEFRLMLCYIYDYFELTVMFDEIDTSGNMLIDAKEFKKAAQKITEWGVKIEDPDAVFKQIDDNGSGQITFDEFAAWASAHKLDADGDPDNV

ATGGGGTGCATTTCGTCTAAGTCGACTTCGTCGAAGAAGGAAGGCAAGAACGCCGCGGAGCGCAAGGTCGCATGGGAGGGCATTCGCCAGCGTCTACCTCGCCGCAAGACTCCTGAGGACAAGGCACGCCGCATTGAGCTCTTCAAACAGTTCGACAAAAACAACACCGGTAAGCTCAGCCTGGAGGAGGTGTACAATGGCTGTGTCACTGTGCTGCATCTGGATGAGTTCACGACTCGCCTGCGCGACATCGTGAAGCGCGCCTTCAGCAAGGCCAAAGACATGGGCACGAAGGACCGCGGCGCGGGTAGCGACCAGTTTGTGGAGTTCCTTGAGTTCCGCCTGATGCTGTGCTACATCTACGACTACTTTGAGCTGACTGTGATGTTCGACGAGATCGATACTTCTGGCAACATGCTGATCGACGCGAAGGAATTTAAGAAGGCTGCTCAGAAGATCACGGAATGGGGTGTGAAGATTGAAGATCCCGACGCCGTGTTCAAGCAGATCGATGACAACGGCTCTGGCCAGATCACGTTCGACGAGTTTGCTGCGTGGGCGAGCGCGCACAAGCTCGACGCTGATGGCGATCCGGACAATGTGGCTGCGTAA

>CbFCaBP2

MGCISSKSTQSKKEGKTAAERKAAWEGIRHRLPRRKTPEDKARRIELFKKFDKNGTGKLSLEETYEGCVSVLHLDEFTTRLRDIVKRAFNKAKEMGTKDRNAGSDQFVEFLEFRLMLCYIYDYFELTVMFDEIDTSGNMLIDAKEFKKAAPKITEWGVKITDPDAVFREIDDNGSGQITFDEFAAWASAHKLDADGDPDNV

ATGGGGTGCATCTCTTCTAAGTCCACTCAGTCGAAGAAGGAGGGCAAGACTGCTGCGGAGCGCAAAGCAGCGTGGGAGGGCATCCGCCACCGCCTGCCTCGCCGCAAGACTCCCGAAGATAAGGCACGCCGTATCGAGCTCTTTAAGAAGTTTGACAAGAACGGTACCGGCAAGCTCAGCTTGGAGGAGACGTACGAGGGCTGCGTATCTGTACTGCACCTGGATGAGTTCACGACTCGTCTGCGCGACATCGTGAAGCGCGCCTTCAACAAGGCAAAGGAAATGGGAACGAAGGACCGCAATGCTGGCAGCGACCAGTTCGTGGAGTTTCTGGAGTTCCGCCTGATGCTGTGCTACATCTACGACTACTTTGAGCTGACAGTGATGTTCGACGAGATTGATACTTCTGGCAACATGCTGATCGACGCGAAGGAGTTCAAGAAGGCTGCTCCGAAGATCACGGAATGGGGTGTGAAGATTACCGACCCCGATGCCGTGTTCAGGGAGATCGACGACAACGGCTCTGGCCAGATCACGTTCGACGAGTTTGCTGCTTGGGCGAGCGCGCACAAGCTGGACGCCGATGGTGATCCGGACAACGTGGCTGCGTAA

>CbFCaBP1

MGCASSAFSSKSKEGKTAAERKVAWGRIRERLPRKKTPEDKERRIELFKKFDVNGTGKLSFEETYRGCIDILHLDEFTTRVRDIVKRAFNKARDMGTKDRNAGSDQFVEFLEFRLMLCYIYDYFELTVMFDEIDTSGNMLIDAKEFKAAVPKIAEWGLKITDPDAVFRKIDGNGSGQITFDEFAAWAAAHRLDADENVYNTK

ATGGGCTGCGCCAGTTCCGCTTTCTCATCCAAGTCCAAGGAGGGCAAGACTGCTGCGGAGCGCAAGGTCGCGTGGGGGAGGATTCGTGAGCGCCTGCCGCGGAAGAAGACTCCCGAGGACAAGGAGCGTCGCATCGAGCTTTTCAAGAAGTTCGATGTGAACGGGACTGGCAAGCTGTCGTTCGAGGAGACGTACAGGGGCTGCATAGATATTCTGCACCTGGATGAGTTCACGACTCGTGTGCGCGACATCGTGAAGCGCGCCTTCAACAAGGCAAGGGACATGGGAACGAAGGACCGCAATGCTGGCAGCGACCAGTTCGTGGAGTTTCTGGAGTTCCGCCTGATGCTGTGCTACATCTACGACTACTTTGAGCTGACAGTGATGTTCGACGAGATTGATACTTCTGGCAACATGCTGATCGACGCGAAGGAATTCAAGGCTGCCGTCCCGAAGATTGCGGAATGGGGTCTGAAGATCACCGACCCCGATGCCGTGTTCAGGAAGATTGACGGCAACGGCTCTGGCCAGATCACGTTCGACGAGTTTGCTGCGTGGGCTGCTGCGCACCGTCTCGATGCCGATGAGAACGTCTACAACACCAAGTAA

>LpFCaBP1

MGCASSLFSSKSKTEDKTAAERKVAWEKIRERLPRRKTPEDKERRIELFKKFDQNNTNRLSVEEVYEGCVDILHLDEFTTRLRDIVKRAFNKAKEMGTKDRGAGSDKFVEFLEFRLMLCYIYDYFELTVMFDEIDTSGNMLIDAKEFKKAVPRIQEWGVKIEDPDAVFKEIDNNGSGQITFDEFAAWAAAHKLDADENPDNTE

ATGGGCTGTGCTTCTTCCCTGTTTTCCTCCAAGTCCAAGACAGAGGATAAAACGGCTGCGGAACGCAAGGTCGCGTGGGAGGGGATCCGCGAGCGTCTGCCTCGCCGCAAGACCCCTGAGGACAAGGAGCGCCGCATTGAGCTCTTTAAGAAGTTCGATCAGAACAACACCAACAGGCTCACCCTTGAGGAGGTCTACGAGGGCTGCGTAGATATTCTGCACCTGGACGAGTTTACGACTCGTCTGCGTGACATTGTGAAGCGCGCCTTCAACAAGGCCAAGGAGATGGGAACGAAGGATCGTGGCGCAGGCAGCGACAAGTTTGTGGAGTTCCTTGAGTTCCGCCTGATGCTGTGCTACATCTACGACTACTTCGAGCTGACTGTGATGTTCGACGAGATCGACACTTCTGGCAATATGCTGATCGACGCAAAGGAGTTCAAGAAGGCTGTTCCAAGGATCCAGGAATGGGGTGTGAAGATTGAGGACCCCGATGCCGTGTTTAAGGAGATCGACAGCAACGGCTCTGGCCAGATCACGTTTGACGAGTTTGCCGCATGGGCGGCAGCGCACAAGCTTGATGCCGATGAGAACCCTGACAACACAGAATAA

>LpFCaBP2

MGCISSKSTQIGKKECKTAAERKAAWEGIRQRLPRRKTPEDKQRRIELFKKFDKNNTNRLTLEEVYEGCVSILHLDEFTTRLRDIVKRAFNKAKEMGTKDRGAGSDKFVEFLEFRLMLCYIYDYFELTVMFDEIDTSGNMLIDAKEFKKAVPRIQEWGVKIEDPDAVFKEIDSNGSGQMTFDEFAAWASAHKLDADGDPDNMAA

ATGTTCCTCTTTTTTTTTTTTCTGTTGGAAGAGCTGTGTCGTCTAGTTTGGTCGTCCGCCCCCCCGTCTAGTTCCCGCTTATTCCCCTCCCCTGTTTGCCCGCCGTGTGTGGCCTCTCTCTACACTCTCGCCTCCCTCCGCTGTAGTTTTTTTCGCGCGCGCACACAATTCTCTCTCCCCAGAGAGAGCCACTATTCTCACTGTGCAAAACTGGGCTGCATCTCTTCAAAGTCTACTCAGATTGGCGAGAGGGAGTGCAAGACGGCTGCGGAGCGCAAGGCCGCGTGGGAGGGGATTCGCGAACGTCTGCCTCGCCGCAAGACCCCTGAGGACAAGGAGCGCCGCATTGAGCTCTTTAAGAAGTTTGATCAGAACAACACCAACAGGCTCACCCTTGAGGAGGTCTACGAGGGCTGCGTGTCTATTCTGCACCTGGACGAGTTTACGACTCGTCTGCGTGACATTGTGAAGCGCGCCTTCAACAAGGCCAAGGAGATGGGAACGAAGGATCGTGGCGCAGGCAGCGACAAGTTTGTGGAGTTCCTTGAGTTCCGCCTGATGCTGTGCTACATCTACGACTACTTCGAGCTGACTGTGATGTTCGACGAGATCGACACTTCTGGCAATATGCTGATCGACGCAAAGGAGTTCAAGAAGGCTGTTCCAAGGATCCAGGAATGGGGTGTGAAGATTGAGGACCCCGATGCCGTGTTTAAGGAGATCGACAGCAATGGCTCTGGCCAGATGACGTTTGACGAGTTTGCCGCATGGGCGGCAGCGCACAAGCTTGACGCCGATGGCGACCCGGACAACATGGCTGCGTAA

>LepFCaBP2-A

MGCTSSKSTQTGKKEGKTAAERKAAWESIRQRLPRQKTAEDKERRIELFKKFDQNGSGKLTFEETYKGCVSVLHLDEFTTRLRDIVKRAFNKAKDMGTKERNSGSEQFVEFLEFRLMLCYIYDYFELTIMFDEIDTSGNMLIDAKEFKKAAPKIAEWGVKISDPDAVFKEIDDNGSGQITFDEFAAWASAHKLDVDGDPDNVAA

ATGGGGTGCACCTCTTCGAAATCTACTCAGACTGGCAAGAAGGAGGGCAAGACTGCCGCGGAGCGTAAGG

CTGCGTGGGAAAGCATCCGCCAGCGTCTGCCCCGCCAGAAGACCGCCGAAGATAAGGAGCGCCGCATTGA

ACTTTTCAAGAAGTTCGATCAGAACGGTTCAGGCAAGTTGACATTCGAAGAGACGTACAAGGGGTGCGTA

TCTGTTCTGCATCTGGATGAGTTCACGACTCGTCTGCGTGATATCGTGAAGCGCGCCTTCAACAAGGCCA

AGGACATGGGAACGAAGGAGCGCAACTCAGGCAGCGAGCAGTTCGTGGAGTTCCTCGAGTTCCGTCTGAT

GCTGTGCTACATCTACGACTACTTTGAGCTAACCATTATGTTCGATGAGATCGACACTTCTGGTAACATG

CTGATCGACGCGAAGGAGTTCAAGAAGGCTGCTCCGAAGATCGCAGAGTGGGGTGTGAAGATCAGCGACC

CTGATGCCGTGTTCAAGGAGATCGACGACAATGGCTCGGGTCAGATCACGTTTGACGAATTCGCCGCCTG

GGCGAGTGCCCACAAGCTCGACGTCGATGGCGATCCGGACAACGTGGCTGCGTAG

>LepFCaBP2-B

MGCTSSKSTQTGKKEGKTAAERKAAWESIRQRLPRQKTAEDKERRIELFKKFDQNGSGKLTFEETYKGCVSVLHLDEFTTRLRDIVKRAFNKAKDMGTKERNSGSEQFVEFLEFRLMLCYIYDYFELTIMFDEIDTSGNMLIDAKEFKKAAPKIAEWGVKISDPDAVFKEIDDNGSGQITFDEFAAWASAHKLDVDGDPDNAE

ATGGGGTGCACCTCTTCGAAATCTACTCAGACTGGCAAGAAGGAGGGCAAGACTGCCGCGGAGCGTAAGG

CTGCGTGGGAAAGCATCCGCCAGCGTCTGCCCCGCCAGAAGACCGCCGAAGATAAGGAGCGCCGCATTGA

ACTTTTCAAGAAGTTCGATCAGAACGGTTCAGGCAAGTTGACATTCGAAGAGACGTACAAGGGGTGCGTA

TCTGTTCTGCATCTGGATGAGTTCACGACTCGTCTGCGTGATATCGTGAAGCGCGCCTTCAACAAGGCCA

AGGACATGGGAACGAAGGAGCGCAACTCAGGCAGCGAGCAGTTCGTGGAGTTCCTCGAGTTCCGTCTGAT

GCTGTGCTACATCTACGACTACTTTGAGCTAACCATTATGTTCGATGAGATCGACACTTCTGGTAACATG

CTGATCGACGCGAAGGAGTTCAAGAAGGCTGCTCCGAAGATCGCAGAGTGGGGTGTGAAGATCAGCGACC

CTGATGCCGTGTTCAAGGAGATCGACGACAATGGCTCGGGTCAGATCACGTTTGACGAATTCGCCGCCTG

GGCGAGTGCCCACAAGCTCGACGTCGATGGCGATCCGGACAACGCCGAATAA

>LepFCaBP1

MGCASSAFSSKSKKEGKTAAERKAAWESIRQRLPRQKTAEDKERRIELFKKFDQNGSGKLTFEETYKGCVSVLHLDEFTTRLRDIVKRAFNKAKDMGTKERNSGSEQFVEFLEFRLMLCYIYDYFELTIMFDEIDTSGNMLIDAKEFKKAAPKIAEWGVKISDPDAVFKEIDDNGSGQITFDEFAAWASAHKLDVDGDPDNAE

ATGGGCTGCGCTTCTTCCGCATTCTCCTCCAAATCCAAGAAGGAGGGCAAGACTGCCGCGGAGCGTAAGG

CTGCGTGGGAAAGCATCCGCCAGCGTCTGCCCCGCCAGAAGACCGCCGAAGATAAGGAGCGCCGCATTGA

ACTTTTCAAGAAGTTCGATCAGAACGGTTCAGGCAAGTTGACATTCGAAGAGACGTACAAGGGGTGCGTA

TCTGTTCTGCATCTGGATGAGTTCACGACTCGTCTGCGTGATATCGTGAAGCGCGCCTTCAACAAGGCCA

AGGACATGGGAACGAAGGAGCGCAACTCAGGCAGCGAGCAGTTCGTGGAGTTCCTCGAGTTCCGTCTGAT

GCTGTGCTACATCTACGACTACTTTGAGCTAACCATTATGTTCGATGAGATCGACACTTCTGGTAACATG

CTGATCGACGCGAAGGAGTTCAAGAAGGCTGCTCCGAAGATCGCAGAGTGGGGTGTGAAGATCAGCGACC

CTGATGCCGTGTTCAAGGAGATCGACGACAATGGCTCGGGTCAGATCACGTTTGACGAATTCGCCGCCTG

GGCGAGTGCCCACAAGCTCGACGTCGATGGCGATCCGGACAACGCCGAATAA
