## Supplementary file 2 for "Protein subcellular relocalization and function of duplicated flagellar calcium binding protein genes in honey bee trypanosomatid parasite"

***Trypanosoma theileri* FCaBPs**

MGLCGSKNSTSSKEGKSAQDRKVAWERIRQAIPREKTPEAKQRRIDLFKKFDKNNSGKLSYNEVYEGCLNVLKLDEFTSRLRDITKRAFNKAKDMGNKVENKGSEDYVEFLEFRLFLCYVYDYFELTVMFDEIDTSGNMLLDENEFKKAAPKLEEWGAKIDDPSKVFKELDKNGSGAVTFDEFAAWASARKLDVDGDPDNVPGNS

ATGGGTTTATGTGGATCTAAGAACAGCACCTCCAGCAAGGAGGGTAAGTCTGCCCAGGACCGCAAGGTTG

CATGGGAGCGCATCCGGCAGGCAATTCCCCGTGAAAAGACACCTGAGGCGAAGCAGCGACGCATCGACTT

GTTTAAGAAGTTCGACAAAAACAATTCTGGGAAGCTGTCTTATAATGAGGTGTATGAAGGCTGTTTGAAT

GTGCTGAAGCTGGATGAATTCACTTCACGGCTACGGGACATCACGAAGCGTGCGTTCAACAAAGCAAAGG

ACATGGGTAACAAGGTGGAGAACAAGGGTTCTGAGGACTATGTTGAATTCCTGGAGTTCCGTTTGTTTCT

GTGCTACGTGTACGACTACTTTGAGCTGACAGTGATGTTCGATGAAATCGACACCTCCGGCAACATGCTG

CTCGATGAGAATGAGTTCAAGAAGGCCGCGCCGAAGCTTGAGGAGTGGGGTGCTAAGATCGATGATCCCA

GCAAAGTGTTCAAGGAGCTGGACAAAAACGGTTCTGGTGCAGTGACGTTCGACGAATTTGCTGCTTGGGC

CTCTGCGCGCAAGCTGGACGTGGACGGTGATCCGGACAACGTTCCAGGCAATTCTTAG

MGSSCSKKEQIKNGFTKKEGKSAQDRKVAWERIRQAIPREKTPEAKQRRADLFKKFDKNNSGKLSYDEVYDGCLNVLKLDEFTTRLRDITKRAFNKAKDMGNKVENKGSEDFVEFLEFRLFLCYVYDYFELTVMFDEIDTSGNMLIDEKEFKKAVPKLEKWGAKIDDPSKVFKELDKNGSGSVTFDEFAAWASARKLDVDGDPDNVPEAPQVPDVPES

ATGGGGTCATCTTGCTCTAAAAAGGAACAAATAAAGAACGGGTTCACCAAGAAGGAGGGTAAGTCTGCCC

AGGACCGCAAGGTTGCATGGGAGCGCATCCGGCAGGCAATTCCCCGTGAAAAGACACCTGAAGCGAAGCA

GCGACGCGCCGACCTGTTTAAGAAGTTCGACAAAAACAATTCTGGGAAGCTGTCTTATGATGAGGTGTAT

GATGGCTGTTTGAATGTGCTGAAGCTGGATGAATTCACTACACGGCTACGGGACATCACGAAGCGTGCGT

TCAACAAGGCAAAGGACATGGGTAACAAGGTGGAGAACAAGGGTTCTGAGGACTTTGTTGAATTCCTGGA

GTTCCGTTTGTTTCTGTGCTACGTGTACGACTACTTTGAGCTGACAGTGATGTTCGATGAAATCGACACC

TCCGGCAACATGCTGATTGATGAGAAGGAGTTCAAGAAGGCCGTGCCGAAGCTTGAAAAGTGGGGTGCTA

AGATCGATGATCCCAGCAAAGTGTTCAAGGAGCTGGACAAAAACGGTTCTGGCTCAGTGACGTTTGACGA

ATTTGCTGCTTGGGCCTCTGCGCGCAAGCTGGACGTGGACGGTGATCCGGACAACGTTCCAGAGGCACCA

CAGGTTCCAGACGTGCCAGAAAGTTCTTAG

MGFCGSKSSTSSKEGKSSQDRKVAWERIRQVIPREKTPEAKQRRIDLFKKFDKNNSGKLSYDEVYEGCLNVLKLDEFTTRLRDITKRAFNRAKDMGNKVANKGSEDFVEFMEFRLFLCYVYDYFDLMVMFDKIDTSGNMLIDEKEFKKAVPKLEKWGAKIDDPSKVFKELDKNGSGSVTFDEFAAWASARKLDVDGDPDNVPGSS

ATGGGGTTTTGTGGATCAAAGAGCAGTACCTCCAGCAAGGAGGGTAAGTCTTCCCAGGACCGCAAGGTTG

CATGGGAGCGTATCCGGCAGGTAATTCCCCGTGAAAAGACACCTGAGGCGAAGCAGCGACGCATCGACTT

GTTTAAGAAGTTCGACAAAAACAATTCTGGGAAGCTGTCTTATGATGAGGTGTATGAAGGTTGTTTGAAT

GTGCTGAAGCTGGATGAATTCACTACACGGCTACGGGACATCACGAAGCGTGCGTTCAACAGGGCAAAGG

ACATGGGTAACAAGGTGGCGAACAAGGGTTCTGAGGACTTTGTTGAATTCATGGAGTTCCGTTTGTTTCT

GTGCTACGTGTACGACTACTTTGATTTGATGGTGATGTTCGATAAAATCGACACCTCCGGCAACATGCTG

ATTGATGAGAAGGAGTTCAAGAAGGCCGTGCCGAAGCTTGAAAAGTGGGGTGCTAAGATCGATGATCCCA

GC

**LpPFRP5**

MTTAASLFERNKRIYLDEESKLHDIVCDDQFLRTFQNWLDQSISKLDFQYERCEQQLSELQQHVSVPKGSFWNSKAIIEHCNLHMAERRLAKRGKELGGDDDGTGDQHVLLPDLIQLVSALQQTKSHSFITAKQRRFIEECEDELKSVVFEPRDLTFITNALQTKLIDDGNTRGVFGDLTAFQNTIEELSPDIDQCEQLLEASIANGEMGLAEDISKRQLDVYEHILRLITDQYPIISNYYSESRNSDRRRRWAVFRMADRDITAVIESKHRQIEACEEDMLKIQEQTTNYNNDDAQQRKRYEADKAESDQFLQQNKEKQQSVWNRVFALFQELQGCSSELATLAEQRRREVERRLQMEEREAGRRSGHESFLQAAAEHAQKLQDTIDNAAAARDVATALNDFVLDGCDSIAAKYDKQQNALGEMLRLVQQHHFKRFSDYYIAASRYLYRKERRLEQIDEEMRSNDMQRELFSDTLNPQAKEYAEANQRLSLQRHEVSQEVMHVRHKLERAERAVTPTLRSLDFANIAYVHPREIVEKMNLSRWSTMLDYRTMLNKSGEDEAELQREAAAIEEMRAELDAQKATSHSQHLLRGATGVRIVTTVPLGATGNKASSKPDAAATAPSSLTCASSVKKQQPSRFVERVYAMLQRGEEPRAGGAAGGGSSSTAAAPKPSSTAAAAGGALAVLPTASLLSPSSLSTRANNNNNDGPLTAGGSMSPAPPPSHIEGATFQALFNYRARAPDELTFEAGQQIICISRAPEEGWFKGVCNQRTGLFPINYVEPVREAASTT

ATGACAACCGCGGCATCGCTGTTTGAGCGCAATAAGCGCATATACCTCGACGAGGAGTCGAAGCTGCACGATATCGTGTGTGACGATCAGTTTCTGCGCACCTTCCAGAATTGGCTGGACCAGAGTATCTCGAAGCTGGATTTCCAGTACGAGCGATGCGAGCAGCAGTTGTCGGAGTTGCAGCAGCATGTGTCGGTCCCGAAGGGTTCCTTCTGGAACAGCAAGGCGATCATTGAGCACTGCAACCTGCATATGGCAGAGCGACGTCTCGCGAAGCGAGGAAAGGAGCTTGGCGGTGACGACGACGGCACGGGCGACCAGCACGTGCTGTTACCGGACCTGATTCAACTTGTGTCGGCGCTGCAGCAGACCAAGTCGCACTCTTTCATCACGGCGAAGCAGCGCCGCTTCATCGAGGAATGCGAGGATGAGCTGAAGTCGGTCGTGTTCGAGCCACGTGATTTGACGTTCATCACAAATGCACTGCAGACAAAGCTGATTGACGACGGCAATACCCGCGGCGTCTTTGGCGACCTCACCGCGTTCCAGAACACGATTGAGGAGCTGTCGCCTGATATCGACCAGTGCGAGCAGCTGTTGGAGGCGAGTATCGCGAATGGCGAGATGGGCCTCGCCGAGGACATCTCGAAGCGGCAGCTGGACGTCTACGAGCACATTCTGCGCCTCATTACAGATCAGTACCCGATCATTTCTAATTACTACTCCGAATCGCGCAACAGCGACCGCCGGCGACGCTGGGCAGTCTTCCGCATGGCGGATCGCGACATCACAGCCGTGATCGAGTCGAAGCATCGCCAAATCGAGGCATGTGAGGAGGACATGCTGAAGATCCAGGAGCAGACGACAAACTACAACAACGACGACGCGCAGCAGCGCAAGCGCTATGAGGCGGATAAGGCCGAATCGGACCAGTTTCTTCAGCAGAACAAGGAGAAGCAGCAGAGCGTGTGGAATCGCGTCTTTGCGCTCTTCCAGGAGCTGCAGGGCTGCTCGAGTGAGCTGGCGACGCTGGCGGAGCAGCGACGGAGGGAAGTGGAGAGACGGCTGCAGATGGAGGAGCGCGAAGCGGGGCGGCGCAGCGGCCACGAGAGCTTCCTGCAAGCCGCGGCCGAGCACGCACAGAAGCTGCAGGACACCATCGACAACGCGGCTGCCGCGCGCGACGTCGCGACGGCGCTCAATGACTTTGTGCTCGACGGCTGCGACAGCATTGCGGCGAAGTACGACAAGCAGCAGAACGCGTTGGGGGAGATGCTGCGGCTAGTGCAGCAGCACCACTTCAAGCGCTTCTCCGACTACTACATCGCAGCCAGTCGCTACCTCTACCGCAAAGAGCGGCGTCTGGAGCAGATTGACGAGGAGATGCGGTCGAACGACATGCAGCGCGAGCTTTTCTCCGACACGCTGAACCCGCAGGCAAAGGAATACGCGGAGGCGAACCAACGACTGTCGCTGCAACGGCACGAGGTGTCGCAGGAGGTGATGCACGTGCGCCACAAGTTGGAGCGGGCAGAACGGGCCGTCACGCCGACGCTGCGCTCGCTTGACTTCGCCAACATCGCCTACGTCCACCCACGCGAGATCGTGGAGAAGATGAACCTGAGTCGCTGGAGCACCATGCTGGACTACCGCACGATGCTGAACAAGTCTGGCGAGGATGAGGCGGAGTTGCAGCGCGAGGCCGCGGCGATCGAGGAGATGCGGGCGGAGCTTGACGCGCAGAAGGCGACGTCGCACAGCCAGCATCTGCTGCGAGGCGCGACTGGGGTGCGGATTGTCACCACCGTACCGCTTGGCGCCACCGGAAACAAGGCAAGCTCGAAGCCGGATGCTGCGGCGACGGCGCCAAGCTCGCTGACGTGTGCGTCCTCCGTGAAAAAGCAGCAACCATCGCGGTTCGTGGAGCGGGTGTATGCAATGCTGCAACGGGGAGAGGAGCCGCGGGCCGGTGGTGCCGCGGGAGGCGGCTCCTCGTCCACCGCCGCCGCCCCGAAGCCGTCCTCCACGGCCGCGGCAGCTGGCGGCGCGTTGGCGGTGCTTCCCACAGCCTCGTTGCTGTCCCCTTCATCCCTCTCCACCCGGGCGAATAACAACAACAACGACGGCCCGCTGACGGCTGGTGGGTCAATGAGCCCAGCCCCGCCACCGTCGCACATCGAAGGCGCCACCTTCCAGGCCCTCTTTAACTACCGCGCACGCGCGCCAGATGAGCTAACCTTTGAAGCAGGGCAGCAGATCATCTGCATTAGTCGCGCACCGGAGGAGGGGTGGTTTAAGGGTGTGTGCAACCAGCGCACAGGGCTGTTTCCCATCAACTACGTGGAGCCGGTGCGCGAGGCGGCAAGCACGACTTGA

**LpBBS1**

MAQKEKSKGESKEKFWLYAFRDHLANLRAFSNCIETADVSGNGDYQLLVADGSKKLKVFGGTALQRELPLFGVPSAIASFYMSTNDAFNKPVIAVATGPYIFMYRNNKPLYRYMIPAVPIDAQESDIWKKLADGVYTVEDAVAKLESLLDSGVQTSSRTLELLLLDTEEERTDFVTRMSAIPLIQMDVATCMTSIPLETLEAEGTSCLVVGTEACFLYVLGAATMEVSLKVVLPSPPVFLIVAGCFAVDYRIIIACRDGRVYSIKHGHLHSAVIQPDAQPCAVARFGNLIAVATTANTLTYYNLKGKKQQSLFLPCPITNLTTITDPITGEDRGLVVALSNGEIRVLVGTQLLHVSLVYGTVTAMKFCRYGRADGALILVLQNGSLVVELLHRNADLTSSKKVETGPPPEQDVPIPVPFLSSVFTAQTSRERKYGADMYQLFQYDLSQLRLTAAKAYLEMVGSGAVPTELGNVTEENEEVAESSLRMNTVVQGLGPVFKVKVQLQNIGAAPLHAVRVVFCLSDDDMYRMPQQVFTIPTLLPSVPLSCEALVELVEGEVKGNAILVVASEPKSTNPLASTLVDLPEAELIEGL

ATGGCGCAGAAGGAAAAAAGCAAAGGGGAGTCGAAGGAGAAGTTCTGGTTGTACGCCTTCCGCGACCACCTCGCCAACCTGCGCGCTTTTTCGAACTGTATCGAGACGGCCGACGTCAGCGGCAACGGCGACTACCAGCTGCTCGTGGCAGATGGCAGCAAAAAACTAAAGGTCTTCGGCGGCACCGCCCTGCAACGCGAGTTGCCTCTCTTTGGCGTGCCGTCGGCGATCGCCTCCTTTTACATGAGCACCAATGACGCCTTCAACAAGCCAGTAATCGCAGTGGCGACGGGGCCGTACATCTTCATGTACCGCAACAACAAACCTCTCTATCGCTACATGATTCCCGCCGTCCCGATTGACGCGCAGGAGTCGGATATTTGGAAAAAGCTCGCTGACGGCGTCTACACCGTCGAGGACGCCGTGGCGAAGTTGGAGTCGCTGCTCGACTCAGGTGTGCAGACCTCGTCGCGGACGCTGGAGTTGCTGCTGCTAGACACGGAGGAGGAGCGGACTGACTTTGTGACGCGTATGAGCGCCATCCCGCTCATTCAGATGGATGTGGCGACTTGCATGACGTCAATCCCGCTGGAGACACTGGAGGCAGAAGGCACGAGCTGCCTGGTCGTGGGTACCGAGGCGTGCTTTCTCTATGTCTTGGGGGCCGCGACGATGGAGGTGTCGCTGAAGGTGGTGCTGCCGAGCCCGCCGGTGTTTTTAATCGTGGCCGGCTGCTTTGCCGTGGATTACCGCATCATCATCGCATGTCGCGACGGCCGCGTTTACTCCATCAAGCACGGCCACCTGCACAGCGCTGTCATCCAGCCTGACGCGCAGCCTTGCGCCGTGGCTCGCTTCGGCAACTTGATCGCCGTTGCCACCACCGCCAACACGCTCACCTACTACAATCTGAAGGGCAAGAAGCAGCAAAGTCTGTTCCTGCCGTGCCCCATCACGAACTTGACCACCATCACTGACCCGATCACTGGAGAGGACAGGGGCCTCGTCGTCGCCCTCAGCAATGGCGAGATTCGCGTGCTGGTCGGCACGCAGCTGCTGCACGTGAGCCTCGTGTATGGTACGGTGACAGCCATGAAGTTCTGCCGCTACGGCCGCGCGGACGGGGCCCTCATTCTCGTCCTGCAGAACGGCTCGCTCGTCGTCGAGCTGCTGCACCGCAACGCCGACCTCACCTCCAGCAAAAAGGTGGAGACGGGCCCGCCACCGGAGCAGGACGTGCCGATCCCCGTGCCCTTTCTCAGCTCTGTCTTTACGGCGCAGACGTCGCGGGAGCGCAAGTACGGCGCGGACATGTACCAGCTTTTCCAGTACGACCTCTCGCAGCTGCGCTTGACAGCGGCAAAGGCGTATTTGGAGATGGTCGGTAGCGGCGCGGTGCCGACGGAGCTGGGCAACGTGACGGAGGAGAACGAGGAGGTGGCGGAGTCGTCCCTGCGCATGAATACTGTGGTGCAGGGGCTTGGGCCCGTCTTTAAGGTGAAGGTACAGCTGCAGAACATCGGTGCGGCGCCGCTGCATGCCGTCCGGGTGGTCTTCTGCCTCTCCGACGACGATATGTATCGCATGCCGCAGCAAGTTTTCACGATTCCGACGTTGCTGCCCTCCGTTCCGCTGTCATGTGAGGCGCTGGTGGAGTTGGTGGAAGGGGAGGTCAAGGGCAACGCAATTTTGGTCGTCGCCTCAGAGCCGAAGAGCACCAATCCGCTTGCAAGCACGTTGGTAGACCTTCCCGAGGCGGAGCTGATTGAGGGGCTGTAA

**LpTULP**

MNPSTPPRPPNQPPHPHGSAASVRFQRTHVVENTGSPTAATTTDAAVAPVLPCTPPTVPLRHPSSLQKQKKPEELVSNDSASEAEDNPNAVIPAGLTVDAVHRAFGSAADASAPPGAGAGVGVGGGGGGGGGAHVAGGGSPSHGIGSSGGRRPGGIIMPTRKEGNDSVSMYSMSQSVLPLDPRERIYYRPRRHLLQCYVERKKQQGHSIASLFPGGHKSFQFFLEHTNDFVLAAVPRNAKSRAVVEDTSDGIGGRFYAVSKGVGNIVFTVNQQQLDNDSRSFVGKLSRRASGLEMVMFSEGDKATRKEIVVVLLENFEDTSRSSFTVVLPAIDAESGYIRPVEGGEVRFLRDGRGGTSVAGGSRAGIDAAMAESSEDEVEATMSTMTKTAPKFAVVDASATPQVGGTVEAPAMAEGAATSSQKKWRTHSLLAKEYRRDPRSPHIIVLKNKVPQWDGVLRGYKLDFHGRATKASEKNFQLVAASDPEKVVMLFGKQSEDRFALDFRYPLCGLQAAAIATTIMTARKMIK

ATGAACCCTTCCACACCACCGCGGCCGCCCAATCAGCCCCCACACCCGCACGGTAGCGCCGCTAGTGTGCGCTTCCAGCGCACTCACGTTGTCGAGAACACCGGAAGCCCGACTGCGGCCACCACCACCGATGCTGCAGTGGCTCCCGTCCTTCCGTGTACGCCGCCCACCGTTCCTTTACGCCACCCGTCCTCCCTGCAGAAACAAAAAAAGCCGGAGGAACTCGTCAGCAACGACTCAGCCAGCGAGGCCGAGGATAACCCGAACGCCGTCATCCCGGCAGGGCTGACGGTGGACGCGGTGCACCGCGCCTTTGGTAGCGCGGCGGATGCCTCCGCGCCACCTGGCGCTGGCGCTGGCGTTGGCGTTGGCGGTGGCGGTGGCGGTGGCGGTGGCGCGCATGTGGCCGGCGGCGGGTCGCCGTCTCATGGGATTGGCAGTAGCGGCGGCCGTCGGCCTGGTGGGATCATCATGCCGACCCGGAAAGAGGGGAACGACTCGGTCTCCATGTACAGCATGTCGCAGTCCGTCCTGCCACTGGACCCGCGAGAGCGCATCTATTACCGCCCGCGCCGTCACCTGCTCCAGTGCTACGTTGAGCGCAAAAAACAGCAGGGCCACAGCATCGCCTCCCTTTTCCCAGGTGGTCACAAGTCCTTTCAGTTTTTCCTCGAGCACACCAACGACTTTGTGCTCGCTGCCGTCCCACGCAATGCCAAGTCGCGCGCCGTGGTTGAGGACACGTCGGACGGCATCGGTGGCCGTTTCTACGCTGTCTCTAAAGGTGTCGGCAACATCGTCTTCACAGTTAATCAGCAGCAGCTGGACAATGACTCGCGCAGCTTTGTCGGCAAGTTGTCCCGGCGTGCGAGCGGGCTGGAGATGGTGATGTTCAGTGAAGGTGACAAGGCAACACGGAAGGAGATCGTGGTCGTGCTGCTCGAGAACTTCGAGGACACGAGCCGCTCGTCATTTACGGTGGTGCTGCCCGCCATCGACGCAGAGTCGGGCTACATCAGACCGGTGGAGGGTGGCGAGGTGAGGTTCCTGCGCGACGGCCGCGGCGGGACGTCGGTGGCCGGCGGCAGTCGCGCCGGCATCGACGCGGCGATGGCCGAGTCGTCCGAGGACGAGGTAGAAGCAACGATGTCGACGATGACGAAGACGGCACCGAAGTTTGCCGTTGTCGACGCGTCCGCCACGCCGCAGGTCGGTGGGACCGTCGAGGCCCCCGCAATGGCGGAAGGGGCCGCGACGAGCTCACAGAAAAAGTGGCGCACGCACAGCTTGCTGGCGAAGGAGTACCGCCGCGACCCGCGCAGCCCACACATCATCGTCTTGAAGAACAAGGTGCCTCAGTGGGATGGCGTGTTGCGGGGTTACAAGTTAGATTTCCACGGTCGTGCCACGAAGGCGAGCGAGAAGAACTTCCAGCTGGTCGCTGCGTCAGATCCTGAGAAGGTCGTGATGTTGTTTGGTAAGCAGAGCGAGGACCGGTTCGCCTTGGACTTCCGCTACCCGCTTTGTGGTTTGCAGGCTGCCGCGATTGCAACGACCATCATGACTGCTCGTAAGATGATCAAGTAA
