## Supplementary file 3 for "Protein subcellular relocalization and function of duplicated flagellar calcium binding protein genes in honey bee trypanosomatid parasite"

| Primer | Sequence (5’→3’) |
| --- | --- |
| LpFCaBP1-5 | TTTCTAGAATGGGCTGTGCTTCTTCCCTGTTTTCC |
| LpFCaBP1-3 | TTTCTAGATTCTGTGTTGTCAGGGTTCTCATCGG |
| LpFCaBP2-5 | TTTCTAGAATGGGCTGCATCTCTTCAAAGTCTACT |
| LpFCaBP2-3 | TTTCTAGACGCAGCCATGTTGTCCGGGTCGCCAT |
| LpPFRP5-5 | TTTCTAGAATGACAACCGCGGCATCGCTGTTTGA |
| LpPFRP5-3 | TTTCTAGAAGTCGTGCTTGCCGCCTCGCGCACCG |
| 1444F | GCCAAGTCTGTGCAAGAGTTCTAG |
| FCaBP1-N16-Rev | AGCCGTCTTATCCTCTGTCTTGGACTT |
| FCaBP1-N28-Rev | CTGACGGATCTTCTCCCATGCTACCTT |
| FCaBP2-N16-Rev | AGCCGTTTTGCACTCCTTCTTGCCAAT |
| FCaBP2-N28-Rev | TTCGCGAATGCCCTCCCACGCGGCCTT |
| FCaBP-487R | CCTCAATCTTCACACCCCATTCCT |
| FCaBP1-N17-For | AAGGAGTGCAAAACGGCTGCGGAACGC |
| FCaBP1-N29-For | TGGGAGGGCATTCGCGAACGTCTGCCT |
| FCaBP2-N17-For | ACAGAGGATAAGACGGCTGCGGAGCGC |
| FCaBP2-N29-For | TGGGAGAAGATCCGTCAGCGTCTGCCT |
| Lp1N16Sense | CTAGAATGGGCTGTGCTTCTTCCCTGTTTTCCTCCAAGTCCAAGACAGAGGATT |
| Lp1N16Antisense | CTAGAATCCTCTGTCTTGGACTTGGAGGAAAACAGGGAAGAAGCACAGCCCATT |
| Lp2N16Sense | CTAGAATGGGCTGCATCTCTTCAAAGTCTACTCAGATTGGCAAGAAGGAGTGCT |
| Lp2N16Antisense | CTAGAGCACTCCTTCTTGCCAATCTGAGTAGACTTTGAAGAGATGCAGCCCATT |
| TbN24Sense | CTAGAATGGGTTGCTCAGGATCGAAGAATACAACGAACTCCAAGGATGGTGCCGCCAGTAAGGGTGGAAAGGACGGGT |
| TbN24Antisense | CTAGACCCGTCCTTTCCACCCTTACTGGCGGCACCATCCTTGGAGTTCGTTGTATTCTTCGATCCTGAGCAACCCATT |
| TcN24Sense | CTAGAATGGGTGCTTGTGGGTCGAAGGGCTCGACGAGCGACAAGGGGTTGGCGAGCGATAAGGACGGCAAGAACGCCT |
| TcN24Antisense | CTAGAGGCGTTCTTGCCGTCCTTATCGCTCGCCAACCCCTTGTCGCTCGTCGAGCCCTTCGACCCACAAGCACCCATT |
| Ld1N15Sense | CTAGAATGGGCTGCAACGCTACCAAGGCTGCCCGGAAGCCCAGCGAGGACT |
| Ld1N15Antisense | CTAGAGTCCTCGCTGGGCTTCCGGGCAGCCTTGGTAGCGTTGCAGCCCATT |
| Cf1N16Sense | CTAGAATGGGTTGCGCTTCTTCCGCTTTTTCCTCCAAGTCCAAGAAGGAGGGCT |
| Cf1N16Antisense | CTAGAGCCCTCCTTCTTGGACTTGGAGGAAAAAGCGGAAGAAGCGCAACCCATT |
| Cf2N16Sense | CTAGAATGGGGTGCATTTCTTCCAAGTCTACTCAGACCGGCAAGAAGGAGGGCT |
| Cf2N16Antisense | CTAGAGCCCTCCTTCTTGCCGGTCTGAGTAGACTTGGAAGAAATGCACCCCATT |
| Ls1N17Sense | CTAGAATGGGTTGCGCCTCTTCCACATCCTCTTCAAAGGGCGGCAAGAAGGAGGGCT |
| Ls1N17Antisense | CTAGAGCCCTCCTTCTTGCCGCCCTTTGAAGAGGATGTGGAAGAGGCGCAACCCATT |
| Ultra-Rev-FCaBP1 | TTGTTCCATATCCTCTGTCTTGGACTTGGA |
| Ultra-For-FCaBP1 | ACAGAGGATATGGAACAAAAACTCATCTCA |
| 3Flag-5 | CTAGAATGGACTATAAGGACCACGACGGAGACTACAAGGATCATGATATTGATTACAAAGACGATGACGATAAGA |
| 3Flag-3 | AGCTTCTTATCGTCATCGTCTTTGTAATCAATATCATGATCCTTGTAGTCTCCGTCGTGGTCCTTATAGTCCATT |
| BBS1-5-HindIII | TTTAAGCTTATGGCGCAGAAGGAAAAAAGCA |
| BBS1-3-ClaI | TTTTATCGATTTACAGCCCCTCAATCAGCTCC |
| TULP-5-HindIII | TTTAAGCTTATGAACCCTTCCACACCACCGCGGC |
| TULP-3-ClaI | TTTTATCGATTACTTGATCATCTTACGAGCAGTCA |
| LpFCaBP-For | TTGTGAGATGGGAACGAAGGATCG |
| LpFCaBP-Rev | AAACCGATCCTTCGTTCCCATCTC |
| LpFCaBP1 5’UTR-F | GAGATGGCAACGAGCCTATGCAGG |
| LpFCaBP1 5’UTR-R | CTTTTTCATTGTTGCAGCACGATCAGAAGTGTT |
| LpFCaBP1 Hph-F | GCTGCAACAATGAAAAAGCCTGAACTC |
| LpFCaBP2 Hph-R | ACATTTTGTCTATTTCTTTGCCCTCGG |
| LpFCaBP2 3’UTR-F | AAGAAATAGACAAAATGTCATTTTCTTTTTTTACCG |
| LpFCaBP2 3’UTR-R | TTTCGATGCCTTCCTTTTCAGCCT |
| LpFCaBP1 5’UTR-Outer-F | GCTAGTGACGTTGGTTTGCTTCTT |
| LpFCaBP-152R | TTCTTAAAGAGCTCAATGCGGCGC |
| Hyg-159R | GCAGCTATTTACCCGCAGGACATA |
| LpFCaBP2 3’UTR-Outer-R | GGACCGCTTTCACCGATCTCAAAA |
| LpFCaBP-538F | ACGTTTGACGAGTTTGCCGCATGG |
| Hyg-846F | CGTATATGCTCCGCATTGGTCTTG |
| LpSL-F | CTAACGCTATATAAGTATCAGTTTCTGTACTTTATTG |
| FCaBP1-36R | GGACTTGGAGGAAAACAGGG |
| FCaBP2-36R | GCCAATCTGAGTAGACTTTG |
| LpGAPDH-F | TGAACGGCCACCGCATCCTG |
| LpGAPDH-R | GGGCCAGGCAGTTGGTCGTG |
| LpITS2-F | GGGTCTTTTGTGATCGGGATAA |
| LpITS2-R | CAAAAAGATGCCTAACGTGAAGAA |
| AmHsTRPA-F | TAGCGTACATGTGGTGCTGT |
| AmHsTRPA-R | GCTAGGCTCCACGTAATCCA |
